# SIR telomere silencing depends on nuclear envelope lipids and modulates sensitivity to a lysolipid drug

**DOI:** 10.1101/2022.07.08.499406

**Authors:** Maria Laura Sosa Ponce, Mayrene Horta Remedios, Sarah Moradi-Fard, Jennifer A Cobb, Vanina Zaremberg

## Abstract

The nuclear envelope (NE) is important in maintaining genome organization. The role of lipids in the communication between the NE and telomere silencing was investigated, including how changes in lipid composition impact gene expression and overall nuclear architecture. For this purpose, yeast cells were treated with the non-metabolizable lysophosphatidylcholine analog edelfosine, known to accumulate at the perinuclear endoplasmic reticulum. Edelfosine treatment induced NE deformation and disrupted telomere clustering but not anchoring. In addition, the association of Sir4 at telomeres measured by ChIP decreased. RNA-seq analysis showed altered expression of Sir-dependent genes located at sub-telomeric (0-10 kb) regions, which was consistent with Sir4 dispersion. Transcriptomic analysis revealed that two lipid metabolic circuits were activated in response to edelfosine, one mediated by the membrane sensing transcription factors, Spt23/Mga2, and the other by a transcriptional repressor, Opi1. Activation of these combined transcriptional programs resulted in higher levels of unsaturated fatty acids and the formation of nuclear lipid droplets. Interestingly, cells lacking Sir proteins displayed resistance to unsaturated fatty acids and edelfosine, and this phenotype was connected to Rap1.

**GRAPHICAL ABSTRACT:** 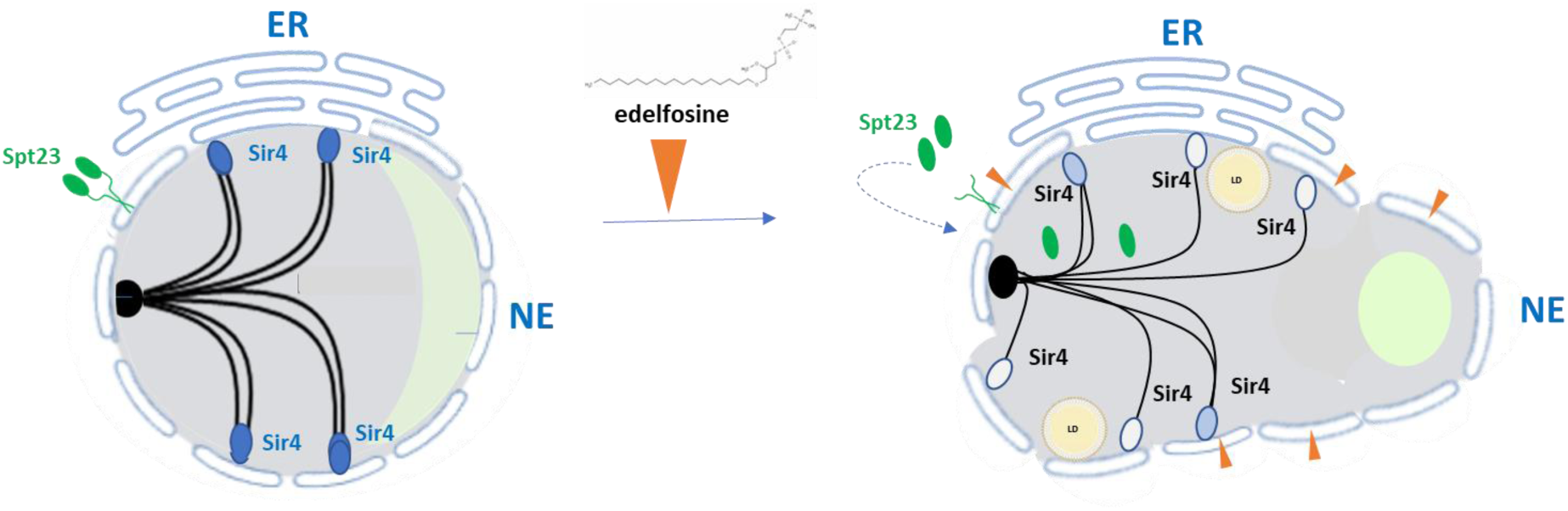

**Summary:** The nuclear envelope (NE) is important for nuclear organization. This study shows that changes in NE lipid composition from lysolipid treatment decreases Sir4 association with telomeres, their clustering at NE, and triggers lipid-specific transcriptional circuits regulated by membrane-sensing factors.

## Introduction

The nuclear envelope (NE) is a double membrane layer contiguous with the endoplasmic reticulum (ER) that protects genomic material from the rest of the cell. The envelope is strategically positioned for sensing stress and transducing signals about lipid homeostasis into the nucleus, and is structurally critical for organizing the genome (Bahmanyar and Schlieker, 2020).

As with all eukaryotes, the nucleus of budding yeast is highly organized. The yeast nucleus is arranged in a Rabl configuration, where centromere clustering at the NE causes chromosomes to occupy spatially constrained territories, promoting clustering between telomeres anchored at the NE (Bystricky et al., 2005; Hozé et al., 2013; Jin et al., 1998). The nucleolus, which is the site of rDNA transcription and ribosome biogenesis, is also tethered to the NE and forms a crescent-shaped subnuclear compartment (Oakes et al., 1993; Taddei and Gasser, 2012; Yang et al., 1989). Therefore, the NE is a key player in the organization of the nucleus.

Interaction with the NE promotes the local accumulation of transcription factors (TFs) and transcription regulators (TRs) at telomeres and rDNA, facilitating silencing (Taddei and Gasser, 2012). The most thoroughly characterized example of proteins that interact with NE-bound anchors involves the formation of heterochromatin by the <u>S</u>ilent Information <u>R</u>egulator (SIR) protein complex, which includes Sir2, Sir3, and Sir4 for deacetylation of lysine 16 of histone H4 (Grunstein and Gasser, 2013). In budding yeast, heterochromatin forms at telomeres and the mating type loci *HML* and *HMR* on chromosome 3. Another silencing factor, Sir1, contributes to transcriptional repression at the silent *HM* mating-type loci, but not at telomeres (Aparicio et al., 1991; Pillus and Rine, 1989). Telomere positioning at the NE occurs through two redundant pathways, where telomere-associated Sir4 binds with the inner nuclear membrane (INM) proteins Esc1 and Mps3, or alternatively where yKu70/80 binds with Mps3 (Hediger et al., 2002; Bupp et al., 2007). Sir4 and Sir3 bind telomeres through interactions with Rap1 (Luo et al., 2002; Moretti et al., 1994), which itself is bound to the double-stranded TG_1-3_ moiety (Buchman et al., 1988a), and together with Sir2, a highly conserved NAD-dependent histone deacetylase (Imai et al., 2000), they form the SIR repressive complex. This complex associates with nucleosomes containing Rap1 bound DNA and self-propagates along sub-telomeric sequences (Grunstein and Gasser, 2013). Telomere anchoring at the periphery increases the relative concentration of SIR proteins, which can also interact in *trans* with SIR proteins bound to other telomeres to form clusters (Gotta et al., 1996; Taddei et al., 2009). The 32 telomeres of a yeast haploid cell typically cluster in ∼3-5 foci during log phase (Laroche et al., 1998; Moradi-Fard et al., 2016). Although anchoring facilitates the formation of repressive compartments at the periphery, which silences sub-telomeric genes, anchoring can occur in the absence of SIR proteins, and association with the nuclear periphery is not a prerequisite for transcriptional silencing (Taddei and Gasser, 2012; Taddei et al., 2009). Importantly, a reduction in SIR protein association with sub-telomeric regions has been observed in response to DNA damage and stress, indicating that regulation of silencing may be a way to control gene expression at specific silent chromatin domains (Mills et al., 1999; Martin et al., 1999; Ai et al., 2002).

As described above, the pathways that regulate NE anchoring and clustering of telomeres depend on reversible interactions between proteins bound to chromosomes and NE membrane proteins. The impact of NE lipids on these pathways and gene expression is currently unknown. However, some links between lipid metabolism and chromatin organization have been previously identified. For example, decreased fatty acid synthesis due to mutations in acetyl-CoA carboxylase coincided with elevated histone acetylation and contribute to the increased expression of factors regulated by histone deacetylases, similar to cells where *SIR2* has been deleted (Galdieri and Vancura, 2012; Papsdorf and Brunet, 2019). Acetyl-CoA derived from the breakdown of fatty acids accounts for a significant portion of the carbon source used in histone acetylation, indicating that transcriptional regulation is impacted by changes in the lipid landscape (McDonnell et al., 2016). Changes in lipid metabolic pathways have also been shown to affect NE size and shape, most notably with the accumulation of phosphatidic acid (PA), a key metabolic intermediate in membrane lipid synthesis, causing a large expansion of the NE, particularly at the nucleolus (Campbell et al., 2006; Witkin et al., 2012; Carman and Han, 2019; Wolinski et al., 2015). The nucleolus-associated NE also defines a membrane subdomain that becomes actively involved in triacylglycerol (TAG) metabolism in response to cell cycle and nutrient signals (Barbosa et al., 2019).

Beyond the nucleolus, not much is known about how other nuclear territories respond to lipid changes at the nuclear envelope. However, there are studies suggesting lipid homeostasis may affect telomere regulation. For example, Sir3 binding and silencing at sub-telomeres is compromised in response to chlorpromazine, an amphiphilic drug that perturbs internal membrane structures by interacting with the polar headgroups of phospholipids (De Filippi et al., 2007; Ai et al., 2002). Additionally, the telomerase antagonist Rif1, a protein that competes with Sir3 and Sir4 for binding to Rap1, is recruited to the NE through palmitoylation (Park et al., 2011; Fontana et al., 2019; Wotton and Shore, 1997). Previous work has also implicated sterols and sphingolipids in the regulation of telomere clustering. Cells lacking Arv1, a factor required for normal intracellular sterol distribution in yeast and mammalian cells (Tinkelenberg et al., 2000), displayed abnormal nuclear morphology and compromised telomere clustering (Ikeda et al., 2015; Papagiannidis et al., 2021). Similar effects were observed when sphingolipid levels decreased in response to pharmacological tools or mutations that inhibited enzymes of the ceramide biosynthetic pathway (Kajiwara et al., 2012; Ikeda et al., 2015). Sterols and sphingolipids are known to form domains in biological membranes based on their preferential interaction, and yeast cells respond to changes in sterol composition of their membranes by adjusting sphingolipids levels (Guan et al., 2009). Lysophosphatidylcholine (lysoPC) lipid analogues disrupt sterol- sphingolipid domains in yeast (Zaremberg et al., 2005). Cells with mutations in ergosterol (*erg3*Δ mutant) or sphingolipid biosynthesis (*lcb1-100* mutant) were hypersensitive to lysoPC related drugs (Zaremberg et al., 2005), while cells lacking chromatin modifiers like Sir4 displayed resistance to these lysoPC analogues (Cuesta-Marbán et al., 2013), pointing to a possible link between membrane lipid alterations and nuclear organization and gene silencing.

We investigated how alterations in membrane lipid composition imposed by a lysoPC burden impact nuclear architecture and gene expression. With this intention, yeast cells were treated with edelfosine, a non-metabolizable lysoPC analogue that accumulates at the NE. Edelfosine belongs to a group of antineoplastics and antiprotozoals known to target membranes. We previously validated budding yeast as a model organism to study this family of non- metabolizable lysoPC lipid analogues using edelfosine as the prototype (Zaremberg et al., 2005). This antitumor therapeutic was shown to function through a novel mechanism involving selective disruption of lipid rafts at the plasma membrane, which in yeast was characterized by sterol internalization and degradation of the lipid raft-associated proton pump Pma1 (Zaremberg et al., 2005; Czyz et al., 2013). Once internalized, edelfosine accumulated at the perinuclear ER (pER) in both yeast (Cuesta-Marbán et al., 2013) and mammalian cells (Bonilla et al., 2015; Gajate et al., 2012). Intriguingly, cells lacking components of the SIR complex like Sir4 displayed resistance to edelfosine (Cuesta-Marbán et al., 2013), opening the possibility that lipid composition in the ER/NE impacts nuclear architecture.

Here we report that edelfosine led to severe changes in the structure of the NE and nuclear associated compartments. We quantified significant changes in nucleolar morphology and telomere clustering, but not telomere anchoring. Using an unbiased RNA-sequencing approach we showed that edelfosine treatment led to a partial loss of silencing in sub-telomeric regions and increased transcription of genes controlled by Spt23 and Mga2 membrane-sensing transcription factors. The inactive membrane-bound forms of Spt23/Mga2 were cleaved in response to edelfosine, resulting in their translocation from the ER membrane into the nucleus. In addition, a dispersion of the SIR complex from telomeres accompanied the transcriptional changes initiated at the NE in the presence of edelfosine. Our work supports a model where disruption of the NE architecture by a lysoPC drug analogue were sufficient to trigger changes in membrane-associated transcription factors and chromatin regulators, connecting nuclear membrane lipid composition to genome organization and transcriptional regulation.

## Results

### Nuclear envelope deformation results in disruption of nuclear architecture

Due to its large phosphocholine head group and a single long saturated acyl tail (18:0), edelfosine has an “inverted cone shape” geometry, inducing membrane bending and disrupting lipid packing (Bierbaum et al., 1979). In addition, the presence of ether linkages to the *sn-1* and *sn-2* positions of the glycerol backbone render this lysoPC analogue metabolically stable, as it cannot be remodelled (Kny, 1969). Given that edelfosine accumulates at the NE (Cuesta-Marbán et al., 2013), we treated yeast cells with edelfosine to alter NE lipid composition in a controlled manner, so that we could investigate its impact on nuclear architecture and transcriptional activity.

Using live imaging in cells expressing the ER marker Sec63^GFP^, we observed a 3-fold increase in non-spherical nuclei after treatment with 20μM edelfosine for 90 minutes (Figure 1A). These deformations in the NE resulted in diverse nuclear shapes deviating from sphericity, with some resembling those seen in yeast with altered PA homeostasis (Campbell et al., 2006; Witkin et al., 2012; Carman and Han, 2019; Wolinski et al., 2015). Asymmetric NE expansion from high PA levels is known to occur in a region of the NE adjacent to the nucleolus, dubbed a nuclear flare (Witkin et al., 2012). Thus, we next examined if deformation of the NE by edelfosine affected nucleolar morphology. Using the nucleolar marker Nop1^CFP^ in combination with the NE marker Nup49^GFP^, we observed condensation and a significant shrinkage of the nucleoli in edelfosine treated cells (Figure 1B). Interestingly, condensed nucleoli were positioned within a flare-like region of the NE induced by edelfosine.

**Figure 1.**
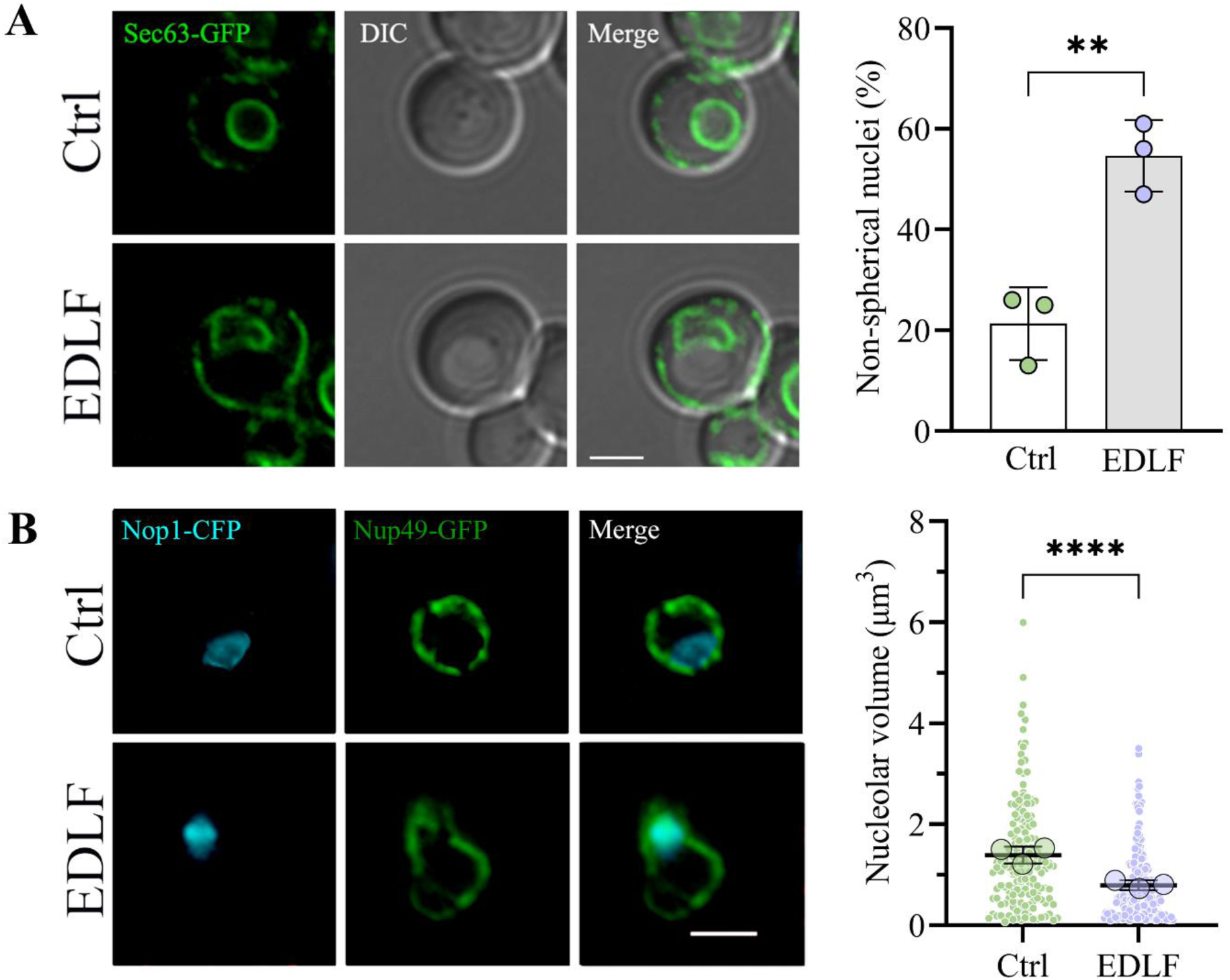
The metabolically stable lysophosphatidylcholine analogue edelfosine alters nuclear envelope morphology and nuclear architecture. (**A**) Representative images of wild- type (W303) cells expressing the ER marker Sec63^GFP^ from a centromeric plasmid. Cells were imaged using live fluorescence microscopy after growth in SD-Leu+Ade and 90 minutes in the presence of edelfosine or the vehicle. Quantification of cells displaying abnormal nuclear membrane morphology is shown beside the microscopy images. Circles represent the percentage of cells displaying abnormal nuclear morphology in each experiment [two-sided Fisher’s exact test gave a p-value < 0.0001 for each experiment (n=100 cells per treatment)]. Bars represent the mean of the three independent experiments ± SD. ** indicates a p-value < 0.01 as determined by two- tailed unpaired t-test (N=3). (**B**) Representative images of wild-type (W303) cells expressing the Nop1^CFP^ nucleolar marker expressed from a centromeric plasmid in cells with Nup49^GFP^ (nuclear envelope marker) endogenously tagged. Cells were imaged using live fluorescence microscopy after growth in SD-Ura+Ade and 90 minutes in the presence of edelfosine or the vehicle. The distribution of values from the quantification of nucleolar volume of cells (described in Materials and Methods) is shown beside the microscopy images. Values cumulative from three independent experiments (n = 163 Ctrl, 190 EDLF) are shown as small circles, with the means of each experiment shown as large circles. Bars represent the mean with 95% confidence interval from the pooled values. **** indicates a p-value < 0.0001 as determined by two-tailed nested t-test (N=3). For all panels: Ctrl = vehicle; EDLF = edelfosine; scale bars represent 2 µm.

Unlike other chemotherapeutic drugs, edelfosine does not target DNA, despite accumulating in the NE. However, edelfosine has been shown to induce DNA fragmentation by production of reactive oxygen species after extended treatment (Renis et al., 2000; Zhang et al., 2007). To determine whether DNA damage was being induced in the timescale of our experiments, the phosphorylation of Rad53 and histone H2A^Ser129^ were analyzed by western blot after edelfosine treatment (Figure S1). No checkpoint signaling response to DNA damage was observed with either of these markers.

To gather further insight about the overall impact of NE deformation, we next investigated other territories and compartments with links to the periphery. Since telomeres show perinuclear anchoring and clustering (Figure 2A), we next determined whether these properties were altered by edelfosine. First, anchoring was assessed by visualizing individual telomeres using the system where lacI-GFP conjugates to lacO arrays integrated in either Tel06R or Tel08L (Figure 2B) (Meister et al., 2010). No significant change in telomere association with the periphery was detected after edelfosine treatment, indicating anchoring continued even with extensive alterations in NE shape, delineated by Nup49^GFP^ (Figure 2C and Figure S2A). We next determined telomere clustering by performing live cell imaging of Rap1^GFP^ foci. In line with previous reports, we observed that 32 telomeres in haploid cells clustered in ∼ 3-5 foci (Figure 2D) (Laroche et al., 1998; Moradi-Fard et al., 2016). Upon edelfosine treatment, this increased to ∼6-7 foci, suggestive of a defect in clustering, where Rap1-bound telomeres are not spatially bundled as effectively (Figure 2D). Consistent with this interpretation, Rap1 recovery by ChIP at three different telomeres, Tel01L, Tel06R, and Tel15L, was statistically similar in the presence and absence of edelfosine (Figure 2E), indicating that the increased number of foci did not result from massive changes in the association of Rap1 with telomeres. Moreover, Rap1 protein levels (Figure 2F) and the fraction of Rap1 associated with the membrane fraction (Figure S2B) did not change, indicating that Rap1 bound telomeres remained anchored at the periphery when the NE was distorted by edelfosine treatment. Importantly, cell cycle analysis by flow cytometry found that cells accumulated in G1 and depleted from S/G2 in response to edelfosine, which indicates that the increase in Rap1 foci is not due to an increase in the number of cells with double DNA content (Figure S2C).

**Figure 2.**
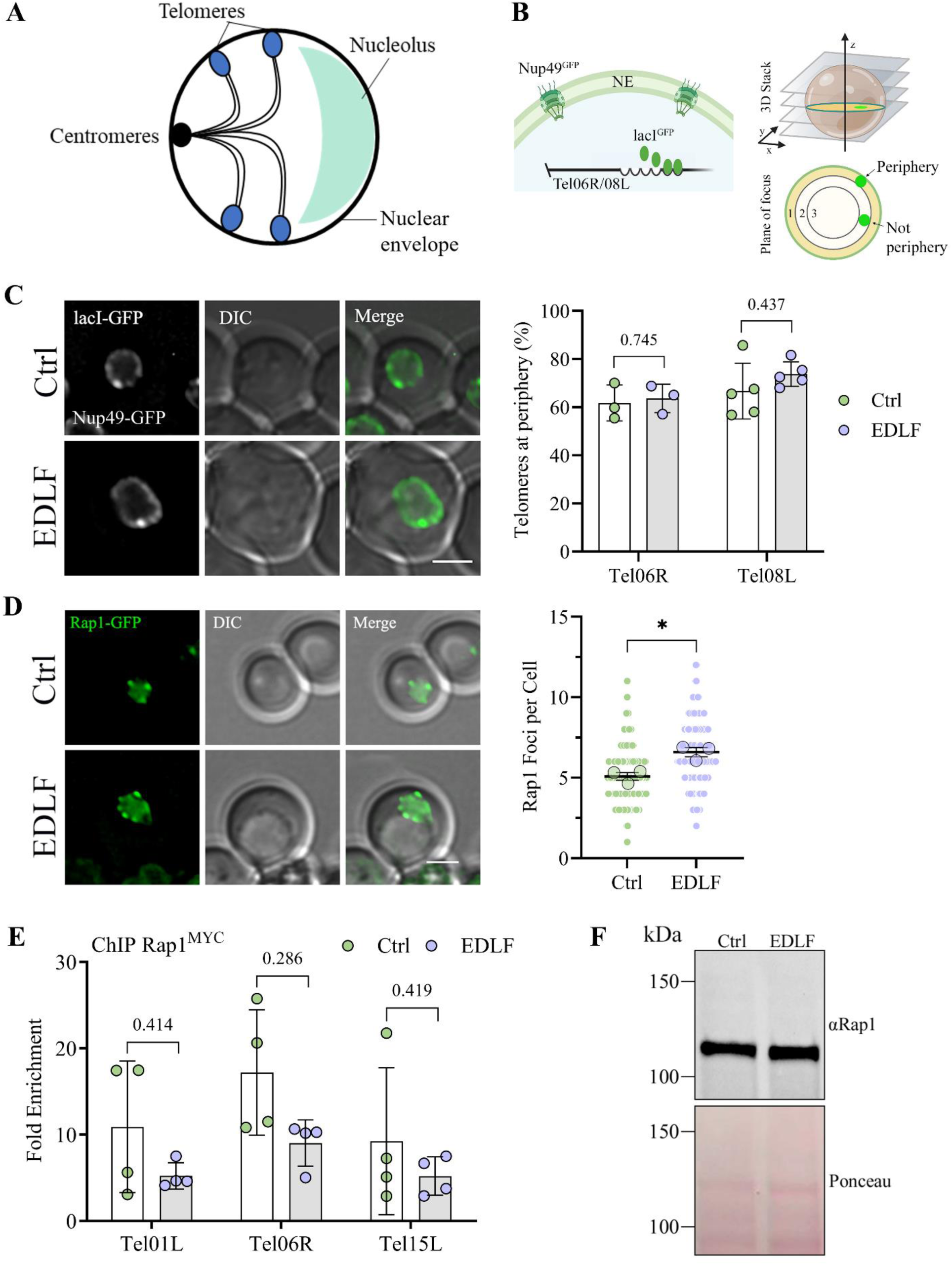
Altered nuclear envelope shape disrupts telomere clustering but not tethering. (**A**) Schematic showing the Rabl conformation of yeast nuclei, where centromeres, telomeres and the nucleolus are tethered to the NE, and telomeres are clustered into 3-5 foci per cell that can be visualized by Rap1. (**B**) Schematic explaining the GFP constructs allowing for analysis of telomere tethering (created with BioRender.com). LacO repeats (6-10 kb) are incorporated into either Tel06R or Tel08L which are then bound by lacI^GFP^ conjugates. The large amount of lacO sites produces a bright focal point at the location of the tagged telomere, which stands out against the nuclear membrane delineated by Nup49^GFP^. (**C**) Representative images of cells expressing the Tel08L constructs described in (B) that were imaged using live fluorescence microscopy after 90 minutes in edelfosine or the vehicle. Quantification of cells showing the telomere at the periphery for both Tel06R and Tel08L is shown beside the microscopy images. Circles represent the percentage of cells displaying telomere localization at the NE in each experiment [two-sided Fisher’s exact test gave the p-values 0.2545, 0.8884 and 0.5461 for each Tel06R experiment (n ≥ 80 cells per treatment), and the p-values 0.0710, 0.1873, 0.3210, 0.6050, and 0.7854 for each Tel08L experiment (n ≥ 45 cells per treatment)]. The bar represents the mean of the experiments ± SD. Differences between control and edelfosine treatments were found to be non-significant by unpaired t-tests with Holm-Šídák correction for multiple comparisons (N=3 for Tel06R, N=5 for Tel08L). (**D**) Representative images of wild-type (W303) cells expressing Rap1^GFP^ imaged by live fluorescence microscopy after 90 minutes in edelfosine or the vehicle. The distribution of values from the quantification of the number of Rap1 foci per cell is shown beside the microscopy images. Values cumulative from three independent experiments (n=180 per treatment) are shown as small circles, with the means of each experiment shown as large circles. Bars represent the mean with 95% confidence interval from the pooled values. * indicates a p-value < 0.05 as determined by two-tailed nested t-test (N=3). (**E**) ChIP-qPCR of Rap1^MYC^ after 60 minutes in edelfosine or control. The fold enrichment at three native sub-telomeres (Tel01L, Tel06R and Tel15L) is shown, normalized to a late replicating region on Chromosome V (469104–469177). Bars represent mean ± SD for four independent experiments while circles represent individual experiments. Differences between control and edelfosine treatments were found to be non-significant by unpaired t-tests with Holm-Šídák correction for multiple comparisons and Welch’s correction for unequal variance (N=4). (**F**) Western blot of endogenous Rap1 in cells expressing 8HIS-Smt3 after 60 minutes with edelfosine or control. Bottom panel shows protein loading by red ponceau. For all panels: Ctrl = vehicle; EDLF = edelfosine; scale bars represent 2 µm.

Altogether, these findings support the notion that NE deformations induced by edelfosine resulted in the loss of NE sphericity affecting all regions of the nuclear membrane, from flares associated with the nucleolus to regions interacting with the bulk of the DNA. These NE deformations, in turn, impacted nuclear architecture, inducing nucleolar compaction and altering telomere clustering, but not anchoring.

### The SIR complex is susceptible to lipid alterations at the NE

We next investigated the SIR complex in edelfosine, as defects in telomere clustering, which can occur independently of anchoring, are associated with dispersion of the SIR complex normally bound at telomeres (Ruault et al., 2011). Moreover, as mentioned above, a chemogenomic screen previously identified *sir4*Δ mutant cells as edelfosine resistant (Czyz et al., 2013; Cuesta-Marbán et al., 2013). Since the screen was performed in the BY4741 background, we first manually verified these results in W303, the background used in the current study. As with *sir4*Δ, deletion of *SIR2* or *SIR3* led to edelfosine resistance compared to wild type (Figure 3A). Expression of *SIR4* from a centromeric plasmid reverted edelfosine resistance in *sir4*Δ (Figure S3A).

**Figure 3.**
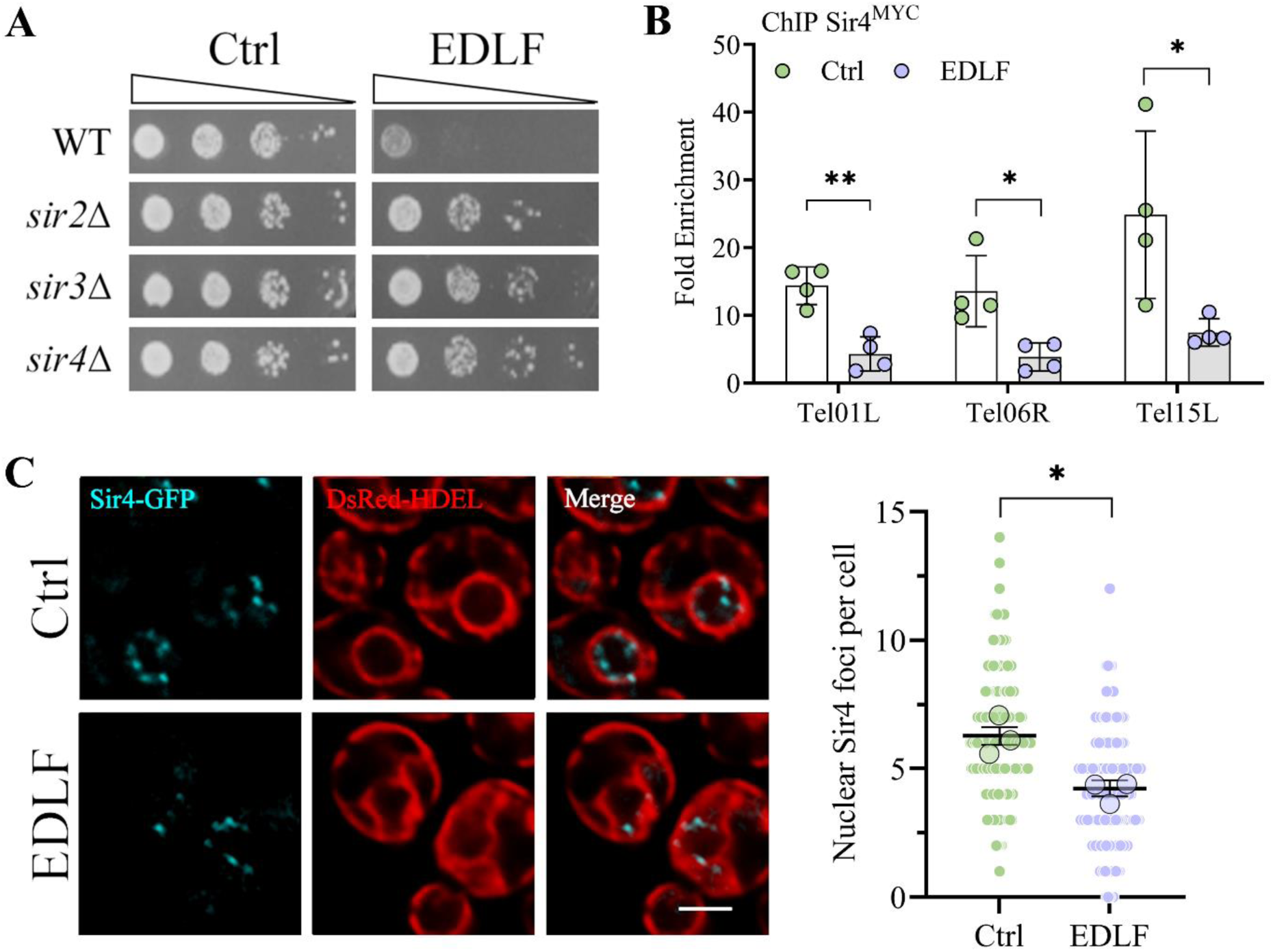
The SIR complex is susceptible to lipid alterations at the NE. (**A**) Wild type (W303) cells or the indicated SIR complex mutants were serial diluted onto synthetic solid media containing 25 μM edelfosine or vehicle and incubated at 30℃ for two days. (**B**) ChIP-qPCR of Sir4^MYC^ after 60 minutes in edelfosine or the vehicle. The fold enrichment at three native sub- telomeres (Tel01L, Tel06R and Tel15L) is shown, normalized to a late replicating region on Chromosome V (469104–469177). Bars represent mean ± SD for four independent experiments while circles represent individual experiments. * indicates p-value <0.05; ** indicates p-value <0.01 as determined by unpaired t-tests with Holm-Šídák correction for multiple comparisons. (**C**) Representative images of cells expressing Sir4^GFP^ and the ER marker ^DsRed^HDEL (both endogenously tagged) visualized by live fluorescence microscopy after 90 minutes with edelfosine or vehicle. Scale bar represents 2 µm. The distribution of values from the quantification of the number of Sir4 foci per cell is shown beside the microscopy images. Values cumulative from three independent experiments (n=180 cells per treatment) are shown as small circles, with the means of each experiment shown as large circles. Bars represent the mean with 95% confidence interval from the pooled values. * indicates a p-value < 0.05 as determined by two-tailed nested t test (N=3). For all panels: Ctrl = vehicle; EDLF = edelfosine.

To explore how edelfosine impacted SIR interactions with telomeres, we next performed ChIP with Sir4^Myc^. There was a marked decrease in Sir4^Myc^ recovery at the three telomeres we monitored (Figure 3B). To complement ChIP, we also performed live cell imaging with Sir4^GFP^ after edelfosine treatment, and observed a reduction in punctate foci formation, which is suggestive of a defect in telomere clustering (Figure 3C) (Moradi-Fard et al., 2016; Palladino et al., 1993). Taken together, these results suggest that deformation of the NE by edelfosine destabilize protein-protein interactions mediated by the SIR complex at telomeres, resulting in Sir4 dispersion and a reduction in telomere clustering. Of note, sensitivity to edelfosine was not affected by C-terminal tagging Sir4 with GFP (Figure S3B).

### Transcriptional changes induced by NE deformation

So far, we have shown that edelfosine induced NE membrane deformations with consequential changes in nuclear architecture, impacting Sir4 association with telomeres and their clustering, as well as nucleolar compaction. These results suggest a loss of Sir-dependent transcriptional regulation at telomeres and decreased transcription in the nucleolus. Yeast cells exposed to various stressors have shown rDNA compaction (reviewed in Matos-Perdomo & Machín, 2019). This phenotype is linked with epigenetic changes as a consequence of altered protein interactions with the NE, as exemplified by rDNA tethering to the NE through the chromosome linkage INM proteins (CLIP) complex (Golam Mostofa et al., 2018; Mekhail et al., 2008; Chan et al., 2011) in combination with the Sir2 histone deacetylase (Gottlieb and Esposito, 1989; Bryk et al., 1997; Fritze et al., 1997; Smith and Boeke, 1997). Similarly, telomere clustering at the NE serves as a platform where protein-protein interactions are strengthened, reinforcing transcriptional silencing (Taddei et al., 2009). Many different NE scaffolding proteins and DNA interacting factors could collaborate to modulate the NE changes imposed by edelfosine into transcriptional changes at telomeres and in the nucleolus. In order to assess the overall transcriptional response, we therefore performed RNA-sequencing (RNA-seq) in the presence of edelfosine. We reasoned that information acquired by this unbiased approach would not only be a resource for understanding the full physiological impact of edelfosine at the molecular level, but also had the potential to reveal unique edelfosine-induced transcriptional changes associated with abnormal NE morphology distinct from the ‘general response’ accompanying most stressors.

A total of 224 genes were found to be differentially expressed in edelfosine when a stringent cut-off of > ln(2)-fold change was used together with a false discovery rate of p < 0.01. Of these, 119 genes were upregulated and 105 were downregulated (complete list in Table S1). We found that 12.6% of the targets upregulated > 2-fold resided in sub-telomeric regions. By contrast, only 4.2% of all genes in the yeast genome are encoded within this same region (p-value 0.0001 by Fisher’s exact test) (Taddei et al., 2009).

To harmonize the observed Sir4 dispersion detected by microscopy and ChIP with sub- telomeric changes in transcription, we queried known SIR targets before moving on to a more expanded analysis of the RNA-seq data. We reasoned that the dispersion of SIRs from telomeres could contribute to the transcriptional changes we detected. Sir2-dependent silencing is prevalent in a zone 0-10 kb from the chromosome ends (Hughes et al., 2000; Bernstein et al., 2000; Ellahi et al., 2015), whereas Hda1 deacetylase acts preferentially within ⁓10-25 kb from telomeres (Robyr et al., 2002), and Rpd3 deacetylase impacts transcription and functions at boundary sites in sub-telomeric regions to prevent the spreading of heterochromatin in opposition to Sir2 (Zhou et al., 2009; Bernstein et al., 2000; Rundlett et al., 1996; Ehrentraut et al., 2010). Of the 120 genes present in the 0-10 kb zone, 29 were upregulated in response to edelfosine when using a cut-off of > ln(0.5)-FC (Table S2).

Two gene families known to be silenced by SIR were represented among this set, including the seri**pau**perin (*PAU*) family (Ai et al., 2002) and the **co**nserved **s**equence (*COS*) gene family (MacDonald et al., 2015; Bedalov et al., 2001). Given that the coding sequence is nearly identical within each family, we designed both specific (*PAU20*, *COS5, COS10*) and general (*PAUG’*, *COSG’*) primers to perform qPCR experiments (Figure S3D, E). As these are known SIR targets, *COS* expression was indeed higher in *sir4*Δ mutants compared to wild type in the absence of edelfosine (Figure S3F). Consistent with the RNA-seq, edelfosine induced a 2.5 ± 0.49 and 1.5 ± 0.82 ln-fold increase in transcription of *PAU* and *COS* genes respectively (Figure S3G). In the cells where *SIR4* was deleted, edelfosine-induced transcription was significantly lower for both gene families compared to wild type, although there was a mild increase in *PAU* expression (1.5 ± 0.23) in *sir4*Δ mutants (Figure S3G).

Furthermore, as expected for Sir4 dispersion considering the protein level of Sir4 remained constant after edelfosine treatment (Figure S3C), known internal (non-sub-telomeric) Sir4 targets represented 22.9% of the downregulated genes in edelfosine (p-value 2.56E-8 by Fisher’s exact test; complete list in Table S3) (Taddei et al., 2009). Taken together, these data support our model that changes in transcription upon edelfosine treatment occur in part through Sir4 disruption and reduced localization to telomeres.

Ribosomal protein genes emerged as the most significantly enriched category of downregulated genes (Figure S4B), consistent with the accompanied rDNA compaction and an expected overall decrease of ribosomal biogenesis. Moreover, in alignment with a recent study demonstrating stress inhibits rRNA processing first rather than rDNA transcription (Szaflarski et al., 2022), we found no significant difference in transcription of the rDNA by qPCR of the 35S fragment (Figure S4C, D).

### Transcription factors mediating changes induced by NE deformation

Next, we looked for over-represented transcription factors (TFs) and transcription regulators (TRs) that could bind to the promoter regions of the two sets of up-and down-regulated genes after edelfosine treatment. Gene promoter analysis with TFs and TRs based on DNA binding was next performed using Yeastract (Monteiro et al., 2020). This analysis identified 6 and 7 TFs/TRs for the up- and down- regulated gene clusters, respectively (p-value <1E-05; Tables 1 and 2). In the cluster of upregulated genes, there was an enrichment of binding sites for TRs involved in the response to oxidative stress and metals. For the cluster of down-regulated genes, we identified Rap1 and its co-regulators of ribosomal protein genes Ifh1 and Fhl1 (Cherel and Thuriaux, 1995; Wade et al., 2004; Rudra et al., 2007), as well as TFs Fkh1 and Fkh2. Interestingly, 36.2% of the down-regulated genes overlapped with known Rap1 targets (Figure 4A, B) (Lieb et al., 2001). Of these, 89.5% were ribosomal protein genes (Figure 4B and Table S4). Representative Rap1 target genes selected for validation by qPCR substantiated the RNA-seq findings (Figure 4C). Taken together with the nucleolar condensation we observed by microscopy (Figure 1B), our data point to a decrease in ribosome biogenesis after edelfosine treatment.

**Figure 4.**
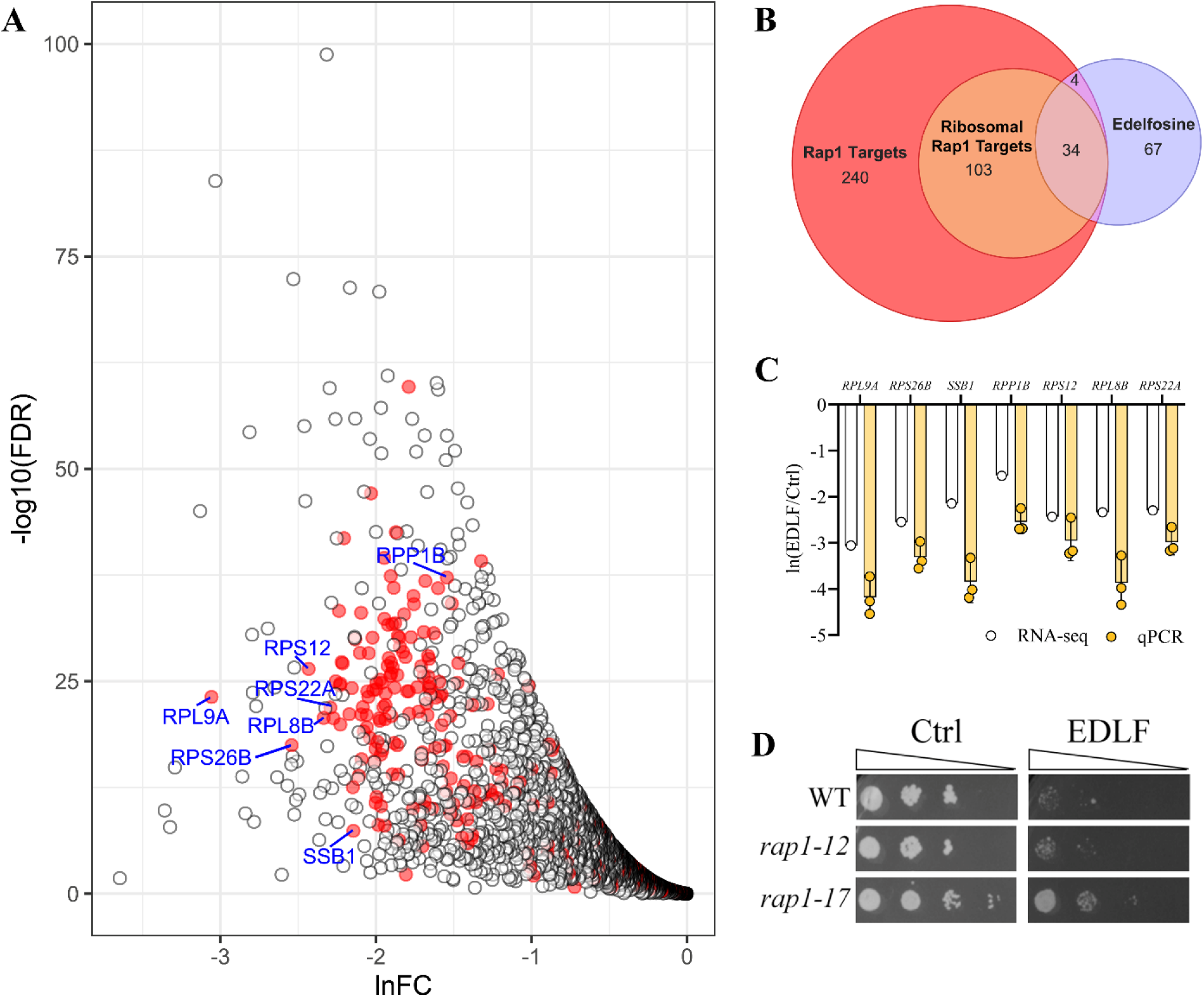
Ribosomal protein targets of Rap1 are repressed in edelfosine. (**A**) Volcano plot of genes downregulated in response to edelfosine as identified by RNA-Seq transcriptome analysis. Targets of Rap1 are coloured red. Genes confirmed by qPCR are labeled. (**B**) Venn diagram showing the overlap between genes strongly downregulated in edelfosine (< -2 ln(FC)) and genes known to be bound by Rap1 (Lieb et al., 2001), including ribosomal protein genes. (**C**) qPCR of genes identified in (A) compared to their RNA-seq values. Bars represent mean ± SD for three independent experiments while circles represent individual experiments. (**D**) The isogenic wild type (KM014) or the indicated Rap1 mutants were serial diluted onto defined solid medium containing 20 μM edelfosine (EDLF) or vehicle (Ctrl) and incubated at 30℃ for two days. The first dilution of *rap1-17* cells was A_600_ ∼ 1, while the other two strains were A_600_ ∼ 0.1, which was experimentally determined to normalize strain growth on control plates.

**Table 1.** Transcription factors with DNA binding sites over-represented in upregulated genes. Analysis of genes with >2-fold increase in response to edelfosine (119 genes) as determined by Yeastract (Monteiro et al., 2020) using DNA binding evidence only. P-values calculated using the hypergeometric test.

| TF | Description | # of genes & (p-value) | References |
| --- | --- | --- | --- |
| **Sko1** | Basic leucine zipper transcription factor of the ATF/CREB family; forms a complex with Tup1p and Cyc8p to both activate and repress transcription; cytosolic and nuclear protein involved in osmotic and oxidative stress responses | 44<br>(4.7E-12) | (Reimand et al., 2010; Niu et al., 2008; Ni et al., 2009) |
| **Hot1** | Transcription factor for glycerol biosynthetic genes; required for the transient induction of glycerol biosynthetic genes GPD1 and GPP2 in response to high osmolarity; targets Hog1p to osmostress responsive promoters; has similarity to Msn1p and Gcr1p | 13<br>(1.2E-09) |  |
| **Spt23** | ER membrane protein involved in regulation of OLE1 transcription; inactive ER form dimerizes and one subunit is then activated by ubiquitin/proteasome-dependent processing followed by nuclear targeting; SPT23 has a paralog, MGA2, that arose from the whole genome duplication | 58<br>(1.3E-08) | (Auld et al., 2006; Reimand et al., 2010) |
| **Yap6** | Basic leucine zipper (bZIP) transcription factor; physically interacts with the Tup1-Cyc8 complex and recruits Tup1p to its targets; overexpression increases sodium and lithium tolerance; computational analysis suggests a role in regulation of expression of genes involved in carbohydrate metabolism; YAP6 has a paralog, CIN5, that arose from the whole genome duplication | 38<br>(6.22E-08) | (Lee et al., 2002; Harbison et al., 2004; Ni et al., 2009; Reimand et al., 2010) |
| **Cad1** | AP-1-like basic leucine zipper (bZIP) transcriptional activator; involved in stress responses, iron metabolism, and pleiotropic drug resistance; controls a set of genes involved in stabilizing proteins; binds consensus sequence TTACTAA; CAD1 has a paralog, YAP1, that arose from the whole genome duplication | 27<br>(1.01E-07) | (Mazzola et al., 2015; Reimand et al., 2010; Haugen et al., 2004; Lee et al., 2002; Azevedo et al., 2007) |
| **Adr1** | Carbon source-responsive zinc-finger transcription factor; required for transcription of the glucose-repressed gene ADH2, of peroxisomal protein genes, and of genes required for ethanol, glycerol, and fatty acid utilization | 31<br>(8.55E-07) | (Chua et al., 2006; Reimand et al., 2010; Karpichev et al., 2008) |

**Table 2.** Transcription factors with DNA binding sites over-represented in downregulated genes. Analysis of genes with >2-fold decrease in response to edelfosine (115 genes) as determined by Yeastract (Monteiro et al., 2020) using DNA binding evidence only. P-values calculated using the hypergeometric test.

| TF | Description | # of genes & (p-value) | Reference |
| --- | --- | --- | --- |
| **Fkh1** | Forkhead family transcription factor with a minor role in the expression of G2/M phase genes; negatively regulates transcriptional elongation; positive role in chromatin silencing at HML and HMR; regulates donor preference during switching | 80<br>(0) | (Pondugula et al., 2009; Knott et al., 2012; Ostrow et al., 2014; Reimand et al., 2010; Workman et al., 2006; Harbison et al., 2004; Lee et al., 2002) |
| **Ifh1** | Coactivator that regulates transcription of ribosomal protein (RP) genes; recruited to RP gene promoters during optimal growth conditions via Fhl1p; subunit of CURI, a complex that coordinates RP production and pre-rRNA processing | 42<br>(0) | (Rudra et al., 2005; Schawalder et al., 2004; Lavoie et al., 2010; Cai et al., 2013; Venters and Pugh, 2009; Wade et al., 2004; Reja et al., 2015) |
| **Spt23** | ER membrane protein involved in regulation of OLE1 transcription; inactive ER form dimerizes and one subunit is then activated by ubiquitin/proteasome-dependent processing followed by nuclear targeting; SPT23 has a paralog, MGA2, that arose from the whole genome duplication | 62<br>(6.6E-14) | (Auld et al., 2006; Reimand et al., 2010) |
| **Fhl1** | Regulator of ribosomal protein transcription; has forkhead associated domain that binds phosphorylated proteins; also has forkhead DNA-binding domain but does not bind DNA in vitro; suppresses RNA pol III and splicing factor prp4 mutants | 53<br>(1.11E-07) | (Rudra et al., 2005; Lee et al., 2002; Tai et al., 2017; Harbison et al., 2004; Kasahara et al., 2007; Lavoie et al., 2010) |
| **Yap6** | Basic leucine zipper (bZIP) transcription factor; physically interacts with the Tup1-Cyc8 complex and recruits Tup1p to its targets; overexpression increases sodium and lithium tolerance; computational analysis suggests a role in regulation of expression of genes involved in carbohydrate metabolism; YAP6 has a paralog, CIN5, that arose from the whole genome duplication | 33<br>(6.0E-07) | (Lee et al., 2002; Harbison et al., 2004; Ni et al., 2009; Reimand et al., 2010) |
| **Fkh2** | Forkhead family transcription factor with a major role in the expression of G2/M phase genes; positively regulates transcriptional elongation; negative role in chromatin silencing at HML and HMR; substrate of the Cdc28p/Clb5p kinase | 53<br>(7.82E-07) | (Pondugula et al., 2009; Harbison et al., 2004; Ostrow et al., 2014; Reimand et al., 2010; Linke et al., 2017; Ghavidel et al., 2018; Gefeng et al., 2000; Simon et al., 2001; Shapira et al., 2004; Workman et al., 2006) |
| **Rap1** | DNA-binding protein involved in either activation or repression of transcription, depending on binding site context; also binds telomere sequences and plays a role in telomeric position effect (silencing) and telomere structure | 59<br>(9.6E-05) | (Chua et al., 2006; Reimand et al., 2010; Lavoie et al., 2010; Workman et al., 2006; Kasahara et al., 2007; Venters and Pugh, 2009; Lieb et al., 2001; De Sanctis et al., 2002; Buchman et al., 1988b; Yarragudi et al., 2007; Lee et al., 2002; Harbison et al., 2004; Hall et al., 2006; Reja et al., 2015) |
| **Ndt80** | Meiosis-specific transcription factor required for exit from pachytene and for full meiotic recombination; activates middle sporulation genes; competes with Sum1p for binding to promoters containing middle sporulation elements (MSE) | 49<br>(1.4E-04) | (Nocedal et al., 2017; Reimand et al., 2010; Chu and Herskowitz, 1998) |

*RAP1* is an essential gene, however a truncated allele missing 165 amino acids at the C-terminus, *rap1-17*, attenuates the repression of ribosomal protein genes in response to secretory pathway defects and disrupts silencing of sub-telomeric regions (Mizuta et al., 1998; Kyrion et al., 1992; Buck and Shore, 1995). By contrast, another characterized allele carrying two missense mutations at amino acids 726 and 727, *rap1-12*, results in loss of silencing at the homothallic mating (*HM*) loci, but does not alter transcription from ribosomal protein genes (Mizuta et al., 1998; Sussel and Shore, 1991; Buck and Shore, 1995). Cell growth on plates containing edelfosine showed that *rap1-17*, but not *rap1-12*, increased edelfosine resistance (Figure 4D). Interestingly, the *rap1-17* C-terminal truncation mutant described above also lacks the domain responsible for interacting with and recruiting Sir4 to telomeres (Cockell et al., 1995; Moretti and Shore, 2001; Luo et al., 2002). In sum, these results support and expand our findings with the SIR complex and suggest the C-terminus of Rap1 facilitates interactions with Sir4 that mediate edelfosine sensitivity.

### Spt23 and Mga2 are activated in response to edelfosine

To understand the mechanism by which changes in the lipid composition at the NE are sensed and communicated to the nucleus, we directed our RNA-seq analysis to differentially expressed genes related to lipid metabolism and transport in response to edelfosine. For these combined categories, an estimated ⁓ 9% (109 genes) and ⁓ 4.5% (60 genes) of the total transcripts differentially expressed [> 0.5 (ln)-fold change cut-off and p < 0.01] were upregulated and downregulated, respectively, in edelfosine. Distinct metabolic branches affecting all lipid classes, including glycerolipids, sphingolipids and sterols, were mapped for each of the up and down datasets (Figure 5A and Table S5). Two critical transcriptional circuits that regulate lipid metabolic pathways clearly emerged from this analysis: Opi1/Ino2/4-mediated repression and Spt23/Mga2 activation. Furthermore, Spt23 was identified as a transcription factor enriched from the RNA-seq analysis (Table 1) and its targets were among the group of transcripts displaying the most dramatic changes in response to edelfosine with >2(ln)-fold change (Figure 5B and Table S5). Spt23, together with its paralog Mga2, maintain NE integrity by way of an ER to nucleus signaling pathway (Zhang et al., 1999). Spt23/Mga2 regulates lipid remodelling by activating expression of *OLE1*, the gene encoding the only fatty acid desaturase present in budding yeast (Zhang et al., 1999; Chellappa et al., 2001; Auld et al., 2006; Martin et al., 2007; Fang et al., 2017; Romero et al., 2018). To confirm activation of the Spt23/Mga2 circuit, we measured transcription of *OLE1* and other targets of Mga2 and Spt23 by qPCR upon edelfosine treatment. Consistent with the RNA-seq results, *OLE1, ICT1, IZH1, IZH2, IZH4* and *MGA2* itself were upregulated while no changes were observed for *SPT23 or IZH3* (Figure 5B, inset). In addition, we also confirmed by qPCR the repression of *INO1*, a representative target of Opi1, and *OPI1* self-repression (Jesch et al., 2005; Nikawa and Kamiuto, 2004) upon edelfosine treatment (Figure 5B, inset).

**Figure 5.**
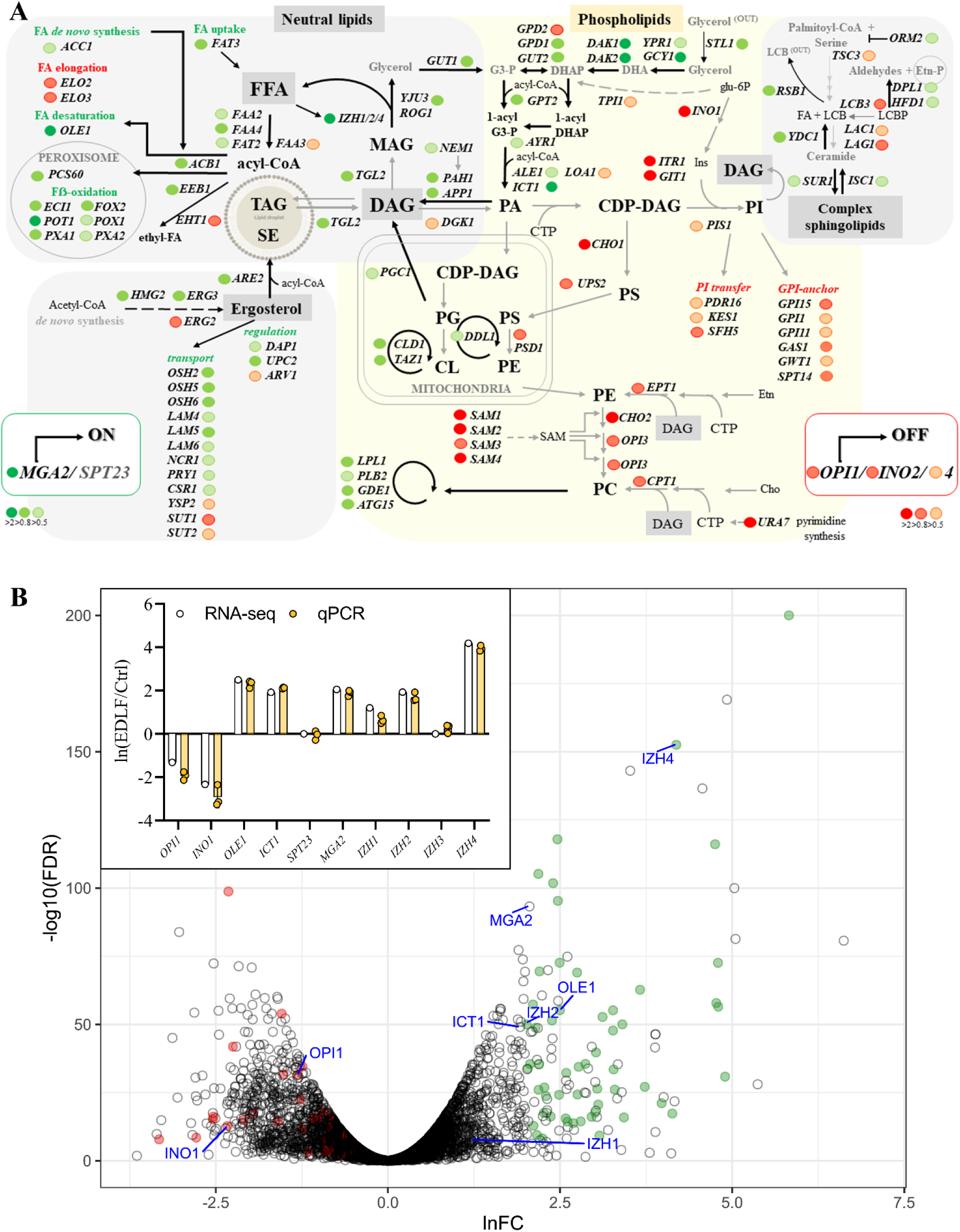
Lipid circuits activated in response to edelfosine. (**A**) Lipid transport and metabolic pathway map highlighting upregulated (green dots) and downregulated (red dots) genes detected by RNA-seq. Color code reflects the fold change as indicated in the figure. (**B**) Volcano plot of transcripts changed in response to edelfosine as identified by the RNA-Seq transcriptome analysis. Targets of Opi1/Ino2/4 are coloured red. Targets of Spt23 are coloured green. Genes confirmed by qPCR (inset) are labeled. For the inset, bars represent mean ± SD for three independent experiments while circles represent individual experiments.

It is worth noting that Spt23 and Mga2 have been previously proposed to be antagonists of silencing (Dula and Holmes, 2000). In fact, 40% of the sub-telomeric genes differentially expressed by treatment with edelfosine can be mapped to the Spt23 regulated network (Table S5). Therefore, Spt23/Mga2 emerged as strong candidates to sense the lipid changes imposed by edelfosine at the NE.

Spt23 and Mga2 form homodimers that insert into the ER membrane by a single transmembrane helix, which is sensitive to the lipid environment (Ballweg et al., 2020). High lipid saturation triggers ubiquitylation, which then signals for the subsequent proteolytic release of a transcriptionally active N-terminal fragment that translocates into the nucleus, inducing *OLE1* expression (Figure 6A) (Hoppe et al., 2000). Based on the enrichment of Spt23 targets in the transcriptomic analysis, we next tested whether edelfosine treatment triggered the processing and subsequent translocation of Spt23 and Mga2 into the nucleus. To this end, we performed live cell imaging of cells expressing either ^GFP^Spt23 or ^GFP^Mga2 from a constitutive promoter wherein proteins were GFP tagged at their N-terminus. Cells harboring either construct showed ER localization (Figure S5), but after a short 30-minute treatment with edelfosine, 98% and 63% of the GFP signal from cells expressing ^GFP^Spt23 and ^GFP^Mga2 respectively were found in the nucleus, indicative of protein processing and translocation (Figure 6B, C). Importantly, the nuclear localization of ^GFP^Spt23 or ^GFP^Mga2 was not a general stress response, as no changes were observed when cells were treated with high levels of methyl methane sulfonate (MMS), an alkylating agent that causes DNA damage. ^GFP^Spt23 and ^GFP^Mga2 processing was further corroborated by western blot. Cleavage of ^GFP^Spt23 could be seen as early as 10 minutes after treatment with edelfosine, with the protein being almost undetectable after 30-minute due to its short half-life (Hoppe et al., 2000) (Figure 6D). ^GFP^Mga2 processing appeared slower (Figure 6E), as both the membrane bound precursor and the soluble active forms were detected after 60 minutes. These results support a role for Spt23 and Mga2 as sensors for changes in the lipid environment imposed by edelfosine accumulation in the NE, and as mediators of the transcriptional response aimed at triggering lipid remodeling in a landscape where the *de novo* synthesis of glycerolipids was repressed.

**Figure 6.**
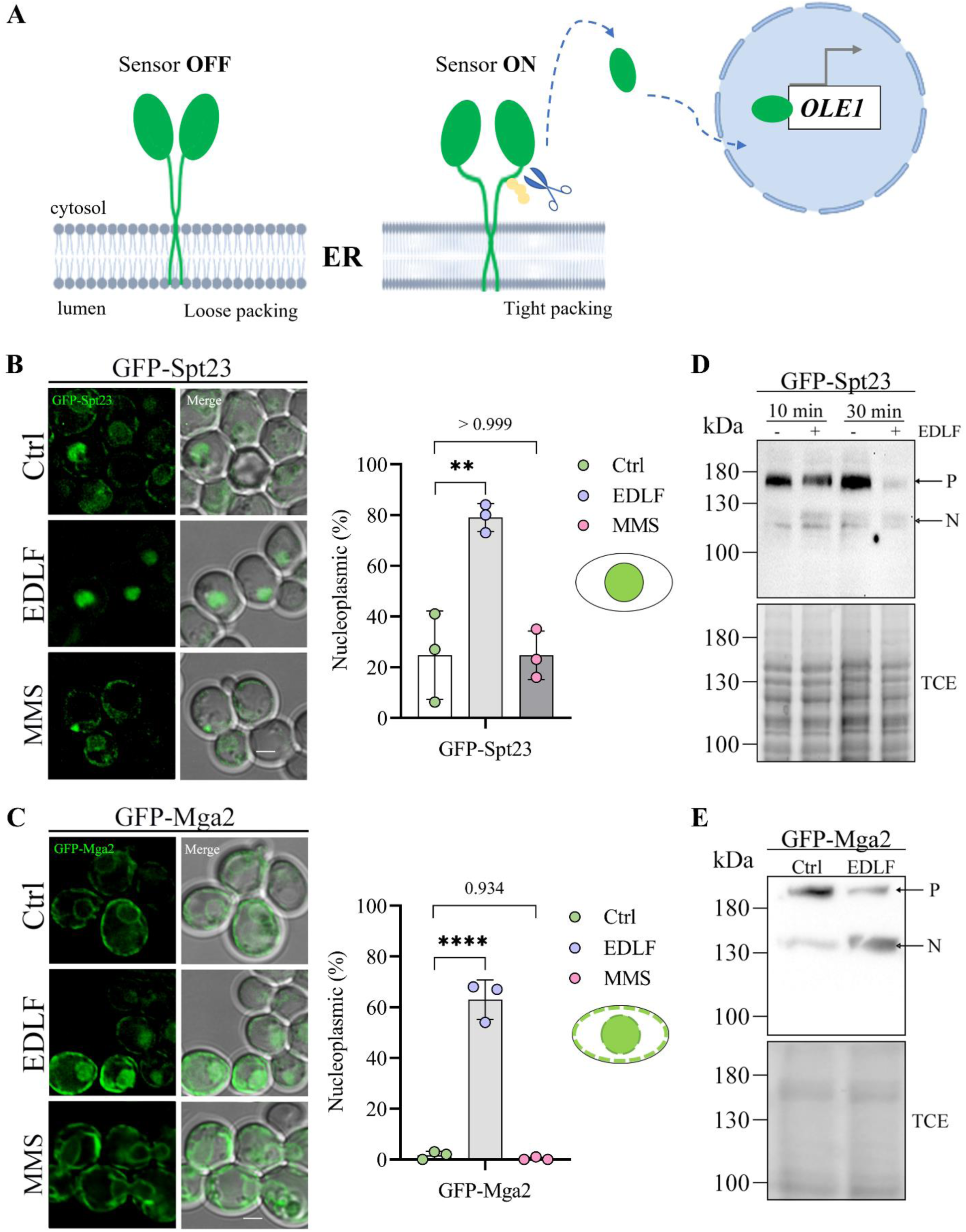
Spt23 and Mga2 sense changes in the membrane lipid environment induced by edelfosine. (**A**) Schematic illustrating the off-on states of the membrane packing sensors Mga2 and Spt23 and the release of the transcription factor N-end in response to changes in membrane environment (created with BioRender.com, adapted from Ballweg et al., 2020). (**B**) Representative images of ^GFP^Spt23 expressed from a centromeric plasmid under the constitutive GPD promoter in *spt23*Δ cells or (**C**) ^GFP^Mga2 expressed from a centromeric plasmid under the constitutive GPD promoter in *mga2*Δ cells were imaged after growth in SD-Leu+Ade and 30 minutes in the presence of edelfosine, the vehicle, or 0.1% methyl methane sulfonate (MMS). Quantification of cells displaying nucleoplasmic signal is shown beside the microscopy images. Nuclear localization was confirmed using the ER marker ^DsRed^HDEL (Supplementary Figure 5). Circles represent the percentage of cells displaying nucleoplasmic signal in each experiment [Chi-square test gave a p- value < 0.0001 for each experiment (n ≥ 65 cells per treatment)]. The bar represents the mean of the three experiments ± SD. ** indicates a p-value < 0.01 and **** indicates a p-value < 0.0001 as determined by standard one-way ANOVA with Tukey’s multiple comparisons test (N=3). Schematics illustrate what was classified as nucleoplasmic signal for B and C. (**D**) Western blot of lysates from cells (W303) expressing ^GFP^Spt23 or (**E**) ^GFP^Mga2 from centromeric plasmids under the constitutive GPD promoter treated with edelfosine for the indicated times (D) or for 60-minutes (E) Bottom panels show protein loading visualized by TCE. P indicates precursor species (148.2kDa for Spt23, 153.9kDa for Mga2), while N indicates nucleoplasmic species (∼90kDa + EGFP). For all panels: Ctrl = vehicle; EDLF = edelfosine; scale bars represent 2 µm.

### Membrane alterations by edelfosine precede changes in nuclear architecture

As mentioned above, Spt23 and Mga2 are believed to function as silencing antagonists (Dula and Holmes, 2000). Edelfosine triggers Spt23/Mga2 processing and subsequent translocation into the nucleus and, as demonstrated above, edelfosine also triggered SIR complex delocalization within the nucleus (Figure 3). The lipid changes associated with NE deformation and Spt23/Mga2 activation could subsequently induce alterations in nuclear organization, including telomere clustering and Sir4 dispersion. Alternatively, a loss of the SIR complex from telomeres in response to edelfosine treatment could result in altered transcription, sensitizing the membrane and resulting in NE conformational changes. To bring insight to these possibilities, we wanted to determine which occurs first: membrane deformation or SIR dispersion. Spt23 and Mga2 are functionally redundant, but the loss of both is lethal (Zhang et al., 1997, 1999), precluding us from determining Sir4 recovery and localization in cells where both were deleted. However, we reasoned that if morphological alterations of the NE and lipid sensing by Spt23 and Mga2 occurred upstream, then changes in nuclear shape and Spt23/Mga2 processing would still occur in the absence of Sir4. Indeed, edelfosine induced NE deformations in *sir4*Δ mutants similarly to *SIR4*^+^cells (Figure 7A). Moreover, ^GFP^Spt23 and ^GFP^Mga2 translocation was similar in *sir4*Δ and wild-type cells upon edelfosine treatment (Figures 7B, C). Interestingly, in *sir4*Δ mutant cells the level of upregulation in some Spt23/Mga2 targets reproducibly decreased after edelfosine treatment, including the transcription of *OLE1* (Figure 7D).

**Figure 7.**
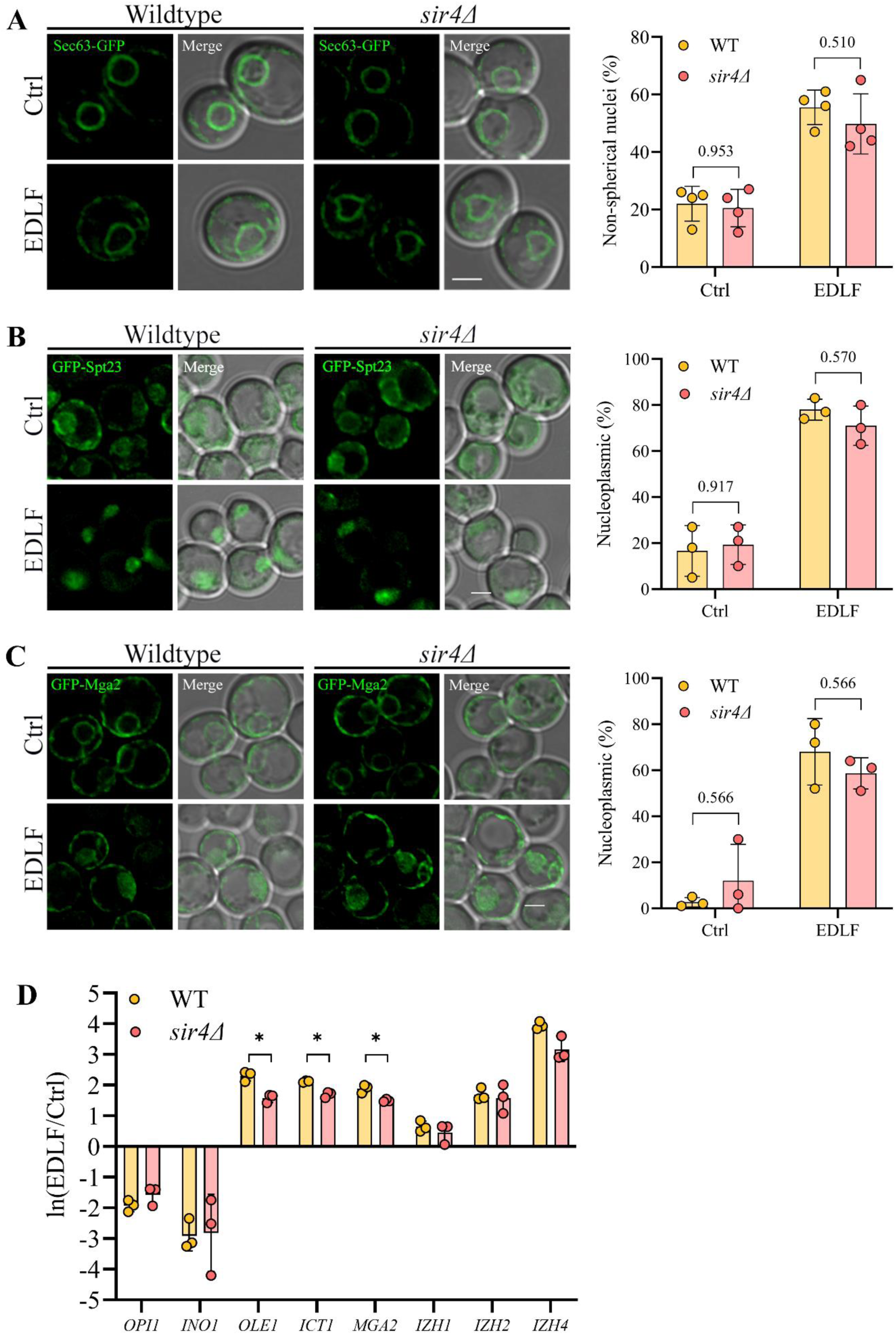
NE deformation and Spt23/Mga2 activation are independent of Sir4. (**A**) Representative images of wild-type and *sir4*Δ cells expressing the ER marker Sec63^GFP^ from a centromeric plasmid. Cells were imaged using live fluorescence microscopy after growth in SD- Leu+Ade and 90 minutes in the presence of edelfosine or control. Quantification of non-round nuclei is shown beside the microscopy images. (**B**) Representative images of ^GFP^Spt23 or (**C**) ^GFP^Mga2 expressed from a centromeric plasmid under the constitutive GPD promoter in wild-type or *sir4*Δ cells imaged after growth in SD-Leu+Ade and 30 minutes in the presence of edelfosine or the vehicle. Quantification of nucleoplasmic signal is shown beside microscopy images. For all microscopy quantifications (**A, B, C**), circles represent the percentage of cells displaying the indicated phenotype in each experiment (n=100 cells per treatment), while the bar represents the mean of all independent experiments ± SD. Differences between wild-type and *sir4Δ* were found to be non-significant in both control and edelfosine treatments as determined by 2-way ANOVA with Šídák’s multiple comparisons test (N=4 for NE, N=3 for Spt23 and Mga2). (**D**) qPCR of Spt23 and Opi1 targets in wild-type and *sir4*Δ cells treated for 60 minutes with edelfosine or vehicle expressed as ln(EDLF/Ctrl). Bars represent mean ± SD for three independent experiments while circles represent individual experiments. * indicates p-value <0.05 as determined by unpaired t tests with Holm-Šídák correction for multiple comparisons. For all panels: Ctrl = vehicle; EDLF = edelfosine; scale bars represent 2 µm.

We next sought to assess if fatty acid unsaturation was augmented in response to edelfosine and if this increase was dependent on Sir4. In line with the upregulation of *OLE1* transcription, neutral lipid analysis of cells treated with edelfosine exhibited higher levels of unsaturated free fatty acids (FFA). Interestingly, this increase was significantly less in *sir4*Δ mutants (Figure 8A). These results support a model whereby edelfosine-induced alterations in lipid composition at the NE are sensed by Spt23/Mga2, resulting in the activation of *OLE1* and a concomitant accumulation of unsaturated FFAs.

**Figure 8.**
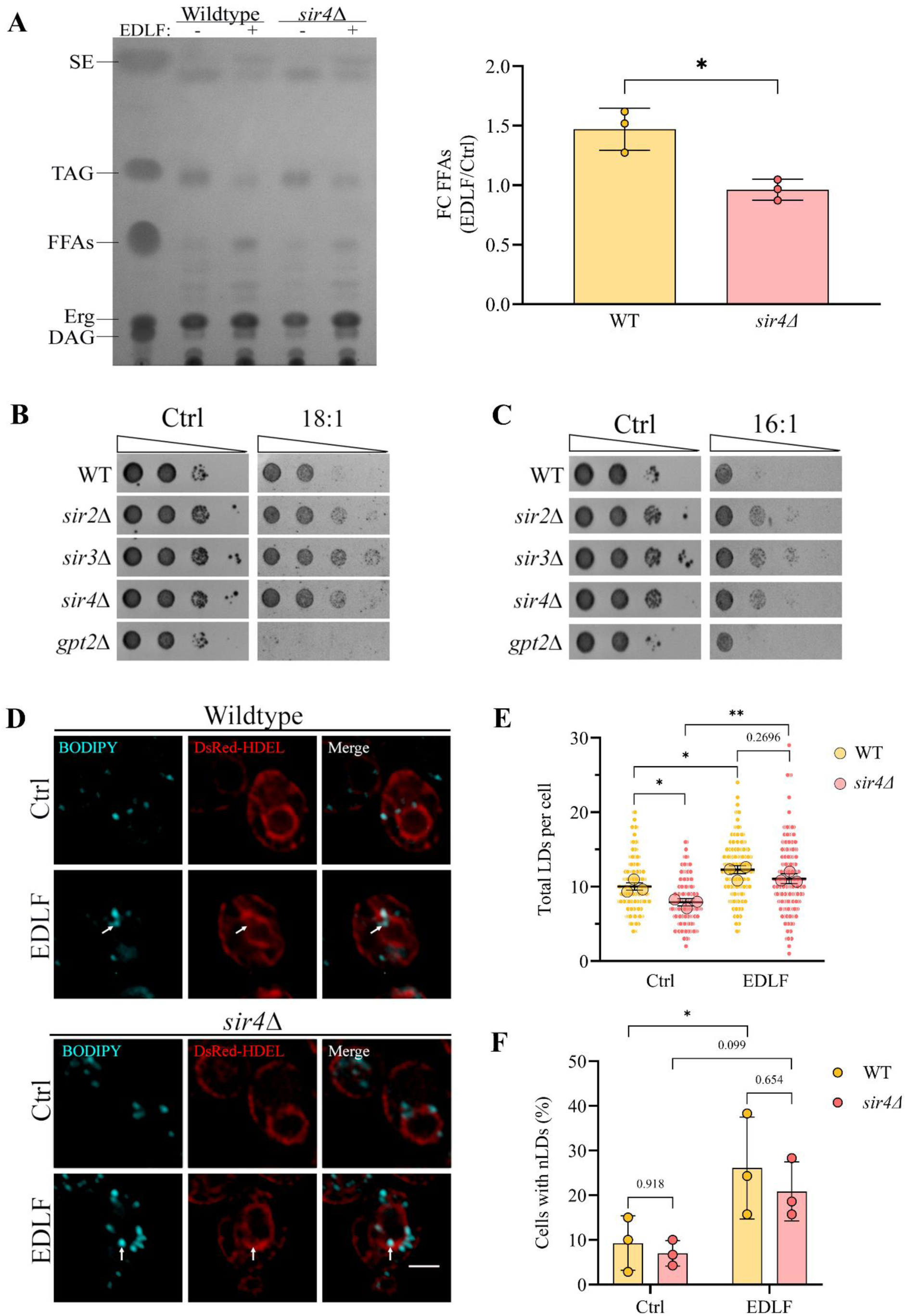
Lipotoxicity response in SIR mutants. (**A**) Wild type or *sir4*Δ cells were treated with edelfosine or the vehicle for 60 minutes, and lipid extractions were performed as described in Materials and Methods. Neutral lipids were separated using thin layer chromatography with a solvent system composed of 80:20:1 petroleum ether/diethyl ether/acetic acid. Diacylglycerol (DAG); ergosterol (Erg); free fatty acids (FFA); triacylglycerol (TAG) and sterol esters (SE). FFA were quantified as described in Methods. The fold-change of total FFA in edelfosine vs control is reported. Circles represent the fold-change for each experiment, while the bar represents the mean of the three experiments ± SD. * indicates a p-value < 0.05 as determined by two-tailed unpaired t-test (N=3). (**B**) Wild type (W303) cells or the indicated mutants were serial diluted onto synthetic solid media containing 4 mM oleic acid or (**C**) 1.18 mM palmitoleic acid or control plates containing glucose and vehicle (1% DMSO). Plates were incubated at 30 ℃ and imaged after three days of growth. (**D**) Representative images of wildtype (BY4741) or *sir4*Δ cells expressing ^DsRed^HDEL treated with edelfosine or the vehicle for 90 minutes and stained with BODIPY^TM^ to visualize lipid droplets (LDs) as described in Materials and Methods. Scale bar represents 2 µm. (**E**) The distribution of values from the quantification of the average number of LDs per cell. Values cumulative from three independent experiments (n ≥ 145 cells per treatment) are shown as small circles, with the means of each experiment shown as large circles. Bars represent the mean with 95% confidence interval from the pooled values. * indicates a p-value < 0.05 and ** indicates a p-values < 0.01 as determined by nested one-way ANOVA with Šídák’s multiple comparisons test (N=3). (**F**) Quantification of cells displaying nuclear lipid droplets (nLDs). Circles represent the percentage of cells displaying nLDs in each experiment (n ≥ 45 cells per treatment)). The bar represents the mean of the three experiments ± SD. * indicates a p-value < 0.05 as determined by 2-way ANOVA with Šídák’s multiple comparisons test (N=3). For all panels: Ctrl = vehicle; EDLF = edelfosine.

A build-up of edelfosine together with unsaturated FFAs could result in a synergistic lipotoxic effect, suggesting that the presence of the SIR complex decreases the lipid detoxifying capacity of yeast. To challenge this idea, we tested the fitness of *sir* mutants on two unsaturated fatty acids commonly found in the yeast lipidome, oleate (18:1) and palmitoleic acid (16:1). Similar to plating on edelfosine, all of the *sir* mutants grew better on these unsaturated FAs compared to wild type (Figure 8B and 8C). The *gpt2*Δ mutant was included as a control as it does not grow on oleate but grows on palmitate (Lockshon et al., 2007; Marr et al., 2012). This suggests that the SIR complex compromises the lipid detoxifying capacity of yeast.

One cellular strategy behind lipid detoxification at the NE is to direct excess fatty acids towards TAG synthesis and lipid droplet (LD) biogenesis (Barbosa et al., 2019; Romanauska and Köhler, 2021). Depending on the status of lipid-related transcriptional circuits, these LDs could accumulate inside the nucleus (nLDs) or face the cytosol (cLDs). When the Opi1/Ino2-4 circuit is repressed by deletion of *INO4,* ∼30% of cells display nLDs in addition to cLDs. By contrast, when *OLE1* is overexpressed, either directly or through constitutive activation of Mga2, only cLDs are produced (Romanauska and Köhler, 2021). Furthermore, when Mga2 is constitutively activated in *ino4*Δ cells, nLDs are still produced. The latter scenario resembles the transcriptional status of cells treated with edelfosine. Since our transcriptomic analysis showed simultaneous Spt23/Mga2 activation and Opi1/Ino2/4 repression, we reasoned the inner nuclear membrane could induce production of nLDs in response to a lysoPC burden. To this end, the number of LDs and their localization was assessed by staining cells expressing the ER marker ^DsRed^HDEL with BODIPY 493/503 (Szymanski et al., 2007) (Figure 8D). In wild type cells, the number of LDs increased from 10.04 ± 3.33 LDs/cell to 12.29 ± 3.61 LDs/cell after edelfosine treatment (Figure 8E). A strong contributor to the increase in LDs was the formation of nLDs inside the nucleus, as ⁓25% of wild type cells treated with edelfosine had at least one nLD (Figure 8F). Although cells lacking Sir4 displayed fewer LDs in control (7.92 ± 3.06 LDs/cell), there was a similar overall increase to 11.07 ± 4.54 LDs/cell upon edelfosine treatment (Figure 8E).

LD bulging from the nuclear membrane towards the nucleoplasm could be a significant contributor to NE deformation, disrupting telomere clustering and overall nuclear architecture independently of transcriptional silencing. Although deacetylation by the SIR complex appears to decrease the lipid detoxifying capacity of yeast since edelfosine and oleate tolerance increases in sir mutant cells, deletion of *SIR4* did not reverse LD formation after edelfosine. In all, these results indicate nLD formation is part of the cellular response to edelfosine, and that it occurs independently of Sir4.

## Discussion

The nucleus, the defining feature of eukaryotic cells, is delimited by a double membrane structure that contains associated integral and peripheral proteins and is vital for the processes regulating nuclear compartmentalization. The architecture of the nucleus is highly regulated, and disruption of its structure has been linked to functional defects such as aberrant gene silencing, loss of chromatin anchoring, incomplete chromosome segregation and faulty cellular differentiation (Golden et al., 2009; Schreiner et al., 2015; Smith et al., 2017; Teixeira et al., 2002). Important, but seemingly often overlooked players in nuclear architecture and regulation, are the lipids that comprise the NE, although perturbations directed at glycerolipid metabolism have been shown to affect NE shape, impacting mainly the NE membrane closely associated with the nucleolus (Santos-Rosa et al., 2005; Siniossoglou et al., 1998; Campbell et al., 2006; Barbosa et al., 2019).

In this study, our work revealed that changing the lipid composition of the NE through a lysolipid burden impacted nuclear architecture and gene expression. Lysolipids are known to introduce membrane deformations and are usually present in small amounts in biological membranes, as they are rapidly re-acylated or metabolized (Fuller and Rand, 2001). Therefore, we manipulated the NE lipid composition by treating cells with edelfosine, a lysoPC analogue that cannot be altered or metabolized. Our data support a model wherein edelfosine-induced changes within the membrane triggered a distinct transcriptional response, indicating contacts between chromatin-associated factors and the nuclear membrane are functionally sensitive to variations in the lipid environment. Deformation of the NE from edelfosine treatment led to changes in genome organization, supporting transcriptional programs aimed at correcting membrane defects and lipid detoxification. Two subnuclear compartments, the nucleolus and telomeres, switched their transcriptional status in response to edelfosine. Telomere clustering decreased and in general transcription of genes in sub-telomeric regions increased, whereas the nucleolus became more compact and there was an overall repression of genes regulating ribosome biogenesis.

### The SIR complex is susceptible to a lysolipid burden

Components of the SIR complex are known to shuttle between telomeres and the nucleolus, the two subnuclear compartments that showed pronounced changes in their transcriptional status during edelfosine treatment (Gotta et al., 1997; Kennedy et al., 1997; Radman-Livaja et al., 2011). SIRs promote telomere clustering, and Sir3 directly promotes long-range contacts between distant chromatin regions, and acts as a molecular bridge between the rDNA and telomeres (Ruault et al., 2021; Hoppe et al., 2002; Radman-Livaja et al., 2011).

SIR association with chromatin at telomeres was highly reactive to an increased burden of lysolipids, like the one imposed by edelfosine. Edelfosine-treated cells led to a reduction in telomere clustering with partial release of Sir4 from telomeres (Figures 2D & 3C) and a reduction in silencing of sub-telomeric regions (Table S2). It is estimated that ∼6% of sub-telomeric genes are silenced by SIRs (Ellahi et al., 2015). While our RNA-seq data showed a loss of silencing at many known SIR targets like *COS7*, *COS4*, *THI5*, *COS8*, *COS1*, *PAU22* and *FDH1* in response to edelfosine (Table S2) (Ellahi et al., 2015; Ai et al., 2002) our data did not perfectly align with all previously reported SIR targets (Ellahi et al., 2015). For example, *IMD2* expression was downregulated 2.8 ln-fold change in response to edelfosine (Table S2), while the opposite effect was seen in sir deletion mutants (Ellahi et al., 2015) suggesting this gene may also be the target of tighter repression by another HDAC or transcription repressor activated by edelfosine. Understanding the overall impact of edelfosine on SIR targets and SIR-independent targets will indeed require additional investigations.

In contrast to Sir4, Rap1 remained bound to telomeres and the telomeres themselves remained anchored after edelfosine treatment. One model to explain these data, which would be in line with how edelfosine functions in lipid bilayers, is that the fluid nature of the membrane decreases where telomeres anchor, impacting the natural dynamics of how they normally cluster (Schober et al., 2008; Hozé et al., 2013). That is to say, while telomere anchoring to the membrane relies on the interaction of chromatin factors with peripheral and integral membrane proteins like Esc1 and Mps3, respectively, clustering relies more on membrane fluidity, allowing the lateral movement of these proteins to bundle and be further stabilized by Sir-mediated *trans* interactions. Edelfosine is known to disturb membrane domain organization at the PM (Ausili et al., 2008; Zaremberg et al., 2005; Cuesta-Marbán et al., 2013; Czyz et al., 2013), and it is reasonable to propose that edelfosine is acting similarly once reaching the ER/NE. An additional contribution to NE deformation and obstruction for telomere clustering may come from the bulging of nuclear LDs in an attempt of cells to recalibrate the fatty acyl composition of the membrane to ease the lysoPC burden.

### Edelfosine-induced repression of ribosomal protein genes

Although Rap1 recovery at telomeres was minimally altered after edelfosine treatment, our study did find that edelfosine induced repression of ribosomal protein genes, which are known to be regulated by Rap1. While a reduction in ribosome biogenesis is part of a general stress response, the signaling pathways leading to a coordinated Rap1-dependent repression of ribosomal protein expression differs depending on the stress. For example, it was previously demonstrated that the *rap1-17* allele prevents ribosomal repression during the secretory stress response, but not the heat shock (Mizuta et al., 1998) or nitrogen starvation responses (Miyoshi et al., 2001). As with the secretory pathway, edelfosine resistance of *rap1-17*, but not *rap1-12,* suggests that repression of ribosomal proteins induced by the lipid drug requires Rap1 C-terminus interaction with Sir4. Altogether, the observed resistance of cells lacking SIR complex components or the C-terminal end of Rap1 supports a model whereby Sir4 binding to Rap1 contributes to edelfosine toxicity (Figures 3A & 4D).

### Sensing lipid changes by Spt23/Mga2 and Opi1, what is being sensed?

Our unbiased RNA-seq approach identified Spt23 and Mga2 transcription factors as membrane sensors that are sensitive to the lipid changes induced by edelfosine. The inactive forms of Spt23/Mga2 reside in the ER/NE membranes and have been previously shown to sense membrane fluidity (Covino et al., 2016), although this concept has been recently challenged by uncovering a particular sensitivity of Mga2 to the degree of lipid saturation instead of viscosity (Ballweg et al., 2020).

What Spt23 and Mga2 are sensing in edelfosine treated cells remains an open question. Activation of these transcription factors was an early event noticed in live cells as early as 30 minutes, thus they might be capable of sensing the saturated C18:0 tail or the change in membrane curvature directly introduced by this lysoPC analogue. Indeed, it is the combination of membrane curvature and composition that modulates lipid packing and protein recruitment (Vanni et al., 2014), opening the possibility that Spt23/Mga2 are sensitive to one or both of these membrane properties.

It is known that edelfosine reduces the stored membrane curvature elastic stress, which is a membrane property that plays a role in the recruitment of several proteins to membranes (Dymond et al., 2008). One such protein is the CTP:phosphocholine cytidylylphosphotransferase (CCT) which localizes to the nucleus and senses curvature through an amphipathic helix (Haider et al., 2018; Taneva et al., 2012; Attard et al., 2000). CCT represents the rate limiting and committed step for the synthesis of PC through the Kennedy pathway, and edelfosine alters its recruitment to membranes while inhibiting the pathway in both yeast and mammalian cells (Boggs et al., 1995; Zaremberg and McMaster, 2002). Of note, it has been shown that in hepatocytes nLDs recruit CCTα (Fujimoto, 2022; Sołtysik et al., 2019). Since we show herein that edelfosine induces nLD biogenesis (Fig 8E), this mechanism could also contribute to the effects previously associated with edelfosine treatment and its impact on CCT activity and the Kennedy pathway.

Opi1 is a transcriptional repressor and the downregulation of Opi1 targets by our RNA-seq analysis suggests that Opi1 is released from the ER and translocated to the nucleus. This occurs in conditions of low PA or in conditions where cytosolic acidification leads to protonation of the phosphate group in PA (Loewen et al., 2004; Young et al., 2010). Edelfosine induces rapid cytosolic acidification (Czyz et al., 2013; Zaremberg et al., 2005). The combined activation of Spt23/Mga2 and Opi1 circuits by edelfosine would result in the upregulation of FA unsaturation and glycerolipid remodelling pathways and a shutdown of de novo glycerolipid synthesis.

### Lipid changes and nuclear territories: insights from edelfosine transcriptomics

The combined analysis of previous lipidomic (Tambellini et al., 2017) and current transcriptomic and neutral lipid profile analysis point to major changes in neutral lipid metabolism leading to unsaturated FA and DAG accumulation (Figure 8A). Importantly, a membrane subdomain linked to lipid droplet biogenesis has been defined at the nuclear envelope tightly connected to the nucleolus (Romanauska and Köhler, 2018). Interestingly, the trans-acylase Lro1 was recently shown to be enriched in this subdomain in response to cell cycle and nutrient signals (Barbosa et al., 2019). Lro1 consumes DAG while generating TAG and LysoPC, which is re-acylated by a PC remodelling pathway (Oelkers et al., 2000). The accumulation of a non-metabolizable LysoPC analogue like edelfosine at the NE would frustrate cellular attempts for re- acylation, resulting in membrane deformation and possible inhibition of TAG synthesis, inducing local DAG accumulation. The lipid metabolic landscape emerging from edelfosine transcriptomic results certainly supports accumulation of DAG through the repression of biosynthetic pathways that consume DAG and the upregulation of several catabolic branches directly producing DAG through TAG lipolysis, phospholipase C, and PA phosphatase Pah1 activities (Figure 5A).

In summary, our work suggests that disruption of the nuclear membrane is sufficient to induce changes in membrane-associated transcription factors and chromatin remodellers. Edelfosine has been studied as a chemotherapeutic, particularly in leukemic bone marrow purging (Vogler and Berdel, 2009; Mollinedo et al., 2004). The current work suggests that the NE could represent a novel target for chemotherapeutics, since lipid drugs like edelfosine are not mutagenic and appear to regulate the activity of sirtuins, which are known to be relevant in the development of multiple chronic diseases (Wang et al., 2012; Bindu et al., 2016; Winnik et al., 2015; Kurylowicz, 2016).

## Materials and Methods

*Reagents.* Unless otherwise indicated, all reagents were purchased from Fisher or Sigma. Yeast extract, peptone, and yeast nitrogen base were from MP Biomedicals (Santa Ana, CA). BODIPY™ 493/503 was ordered from Fisher Invitrogen (D3922). All lipids were ordered from Sigma [1-2- dioleyl-*sn*-glycerol (800811C), oleic acid (O1008-1G), 1,2,3,-trioleylglycerol (870110O) and cholesteryl oleate (C9253)], except for ergosterol (Fluka Analytical, 45480). Edelfosine was the kind gift of Medmark Pharma GmbH.

### Yeast strains, plasmids and primers

Detailed information on yeast strains, plasmids and primers used in this study is provided in Tables 3, 4 and 5 respectively. Strains expressing DsRed-HDEL were transformed with the integrating plasmid pAM41 using the standard transformation protocol (Guthrie and Fink, 1991). Strain JC4881 was created by mating and tetrad dissection, as described previously (Morin et al., 2009). Briefly, parental strains of opposite mating types were streaked on YPD with one intersecting area to allow mating and produce diploids. Diploids were selected and sporulation was then induced by streaking on 2% potassium acetate plates Tetrads were identified by light microscopy and their cell walls were digested with 5 ug/mL Zymolyase (amsbio, 120491-1) in 1M sorbitol for 5 minutes at 30℃. Tetrads were then dissected using a Zeiss Axioskop 40 dissecting microscope and progeny expressing only the markers of interest were saved as the indicated strains. Genotypes were confirmed by colony PCR and sequencing.

**Table 3.**
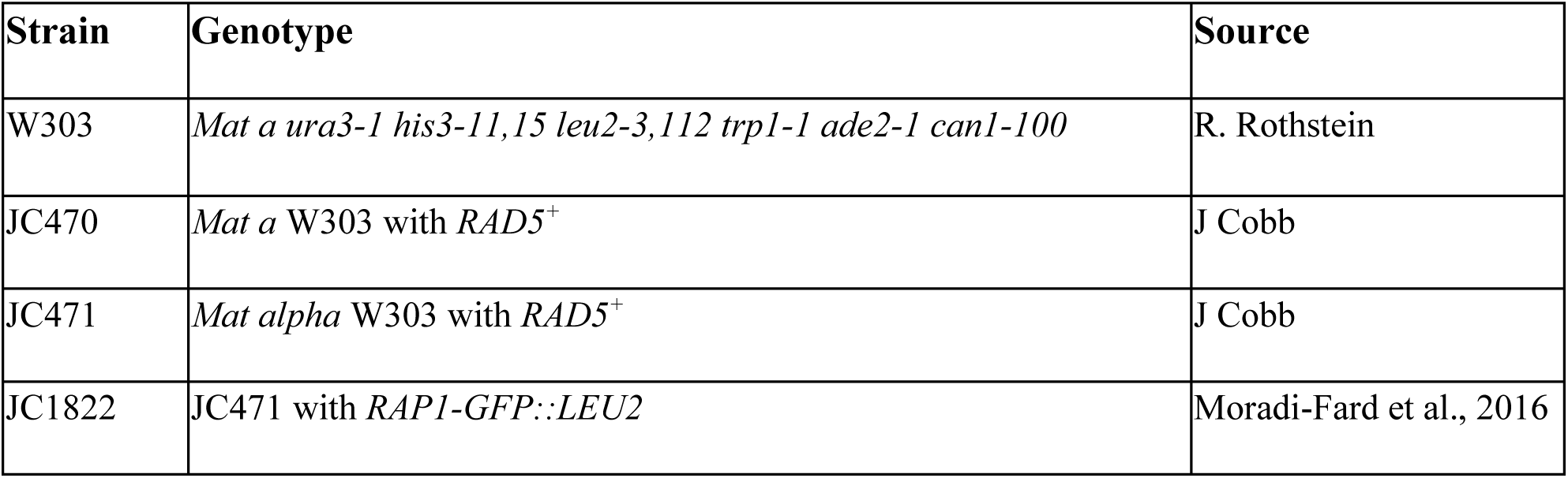
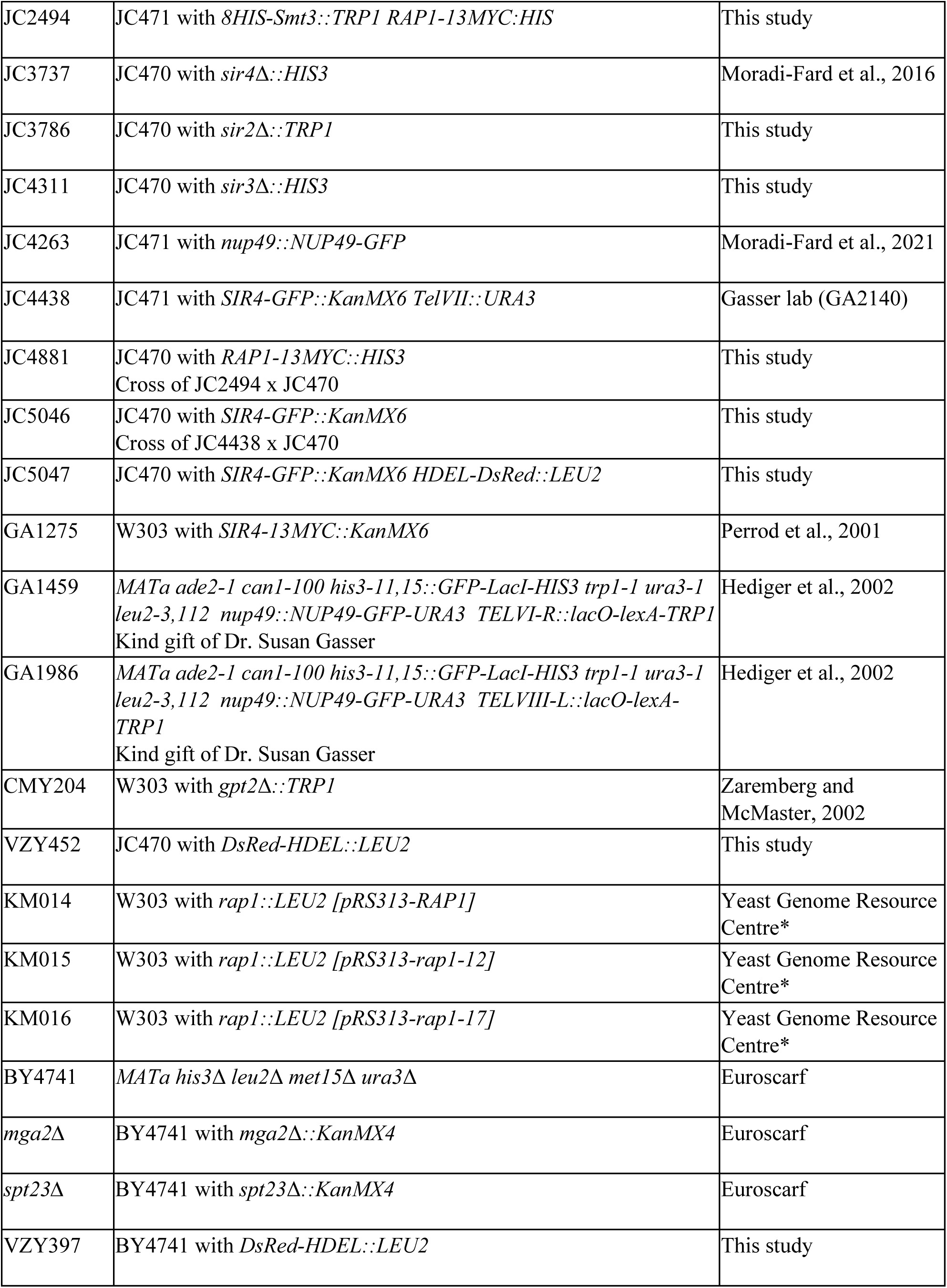
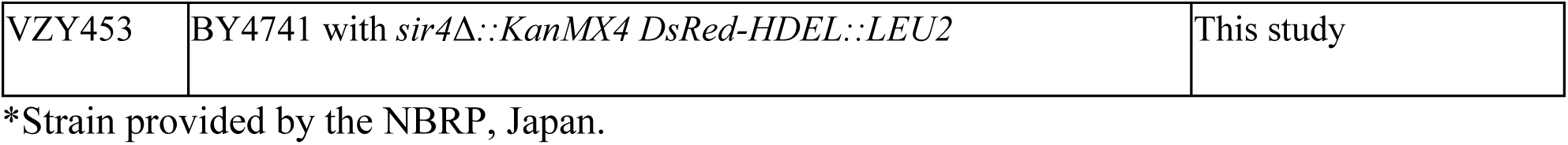
Yeast strains used in this study

**Table 4.**
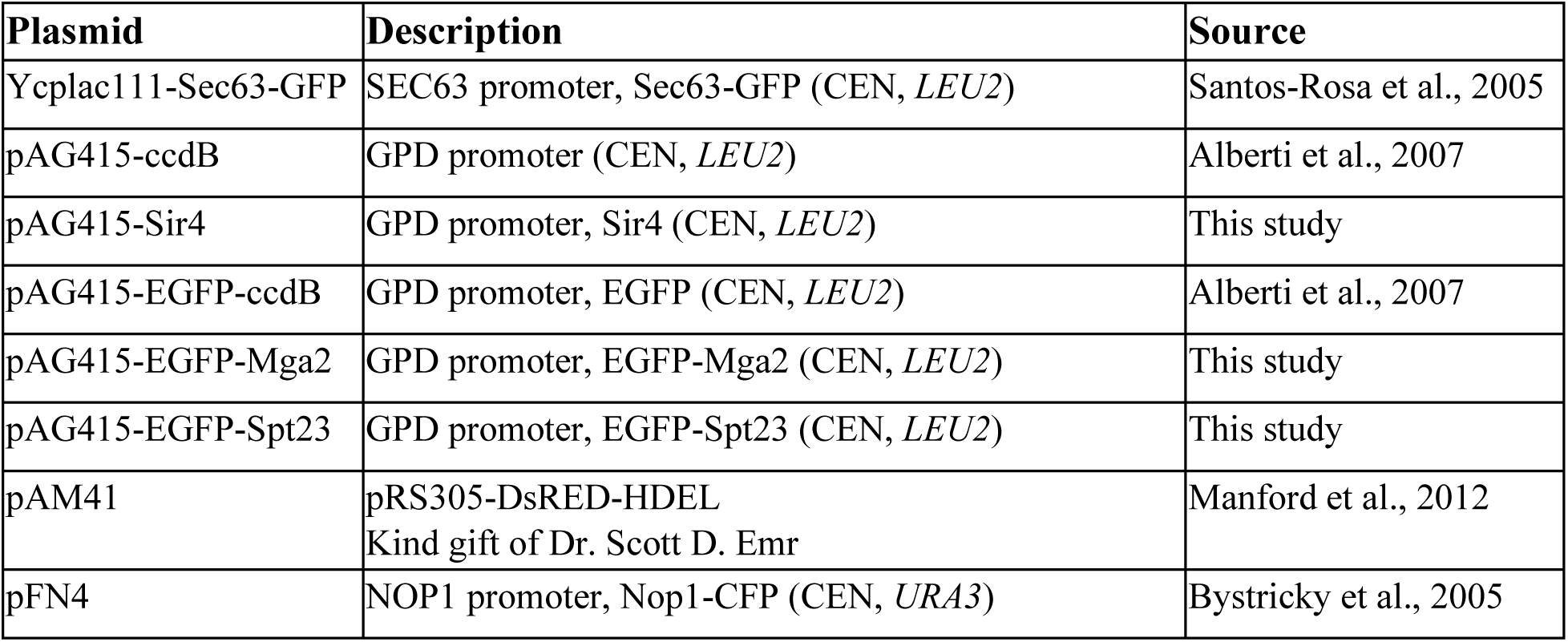
Plasmids used in this study

**Table 5.**
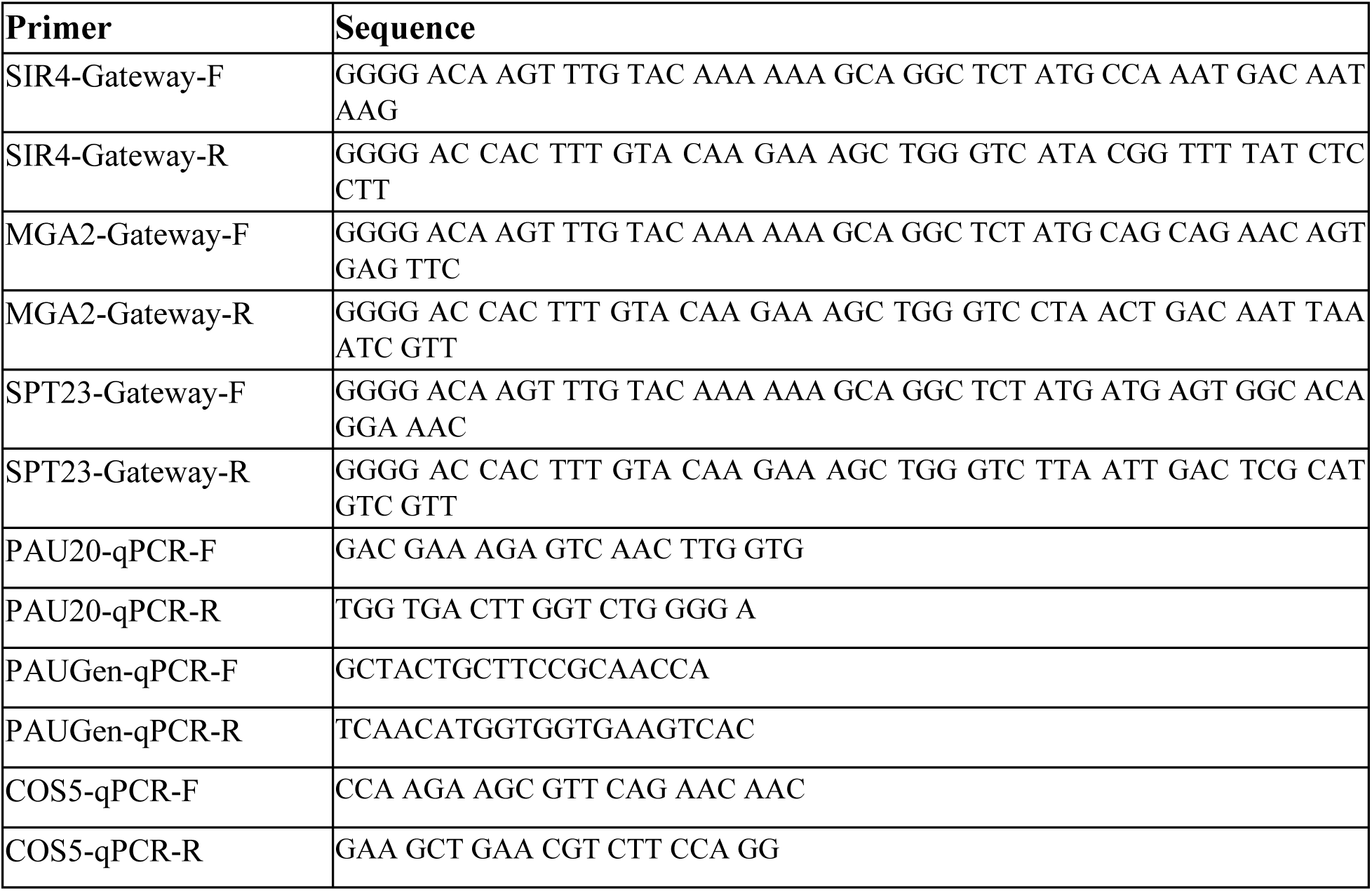
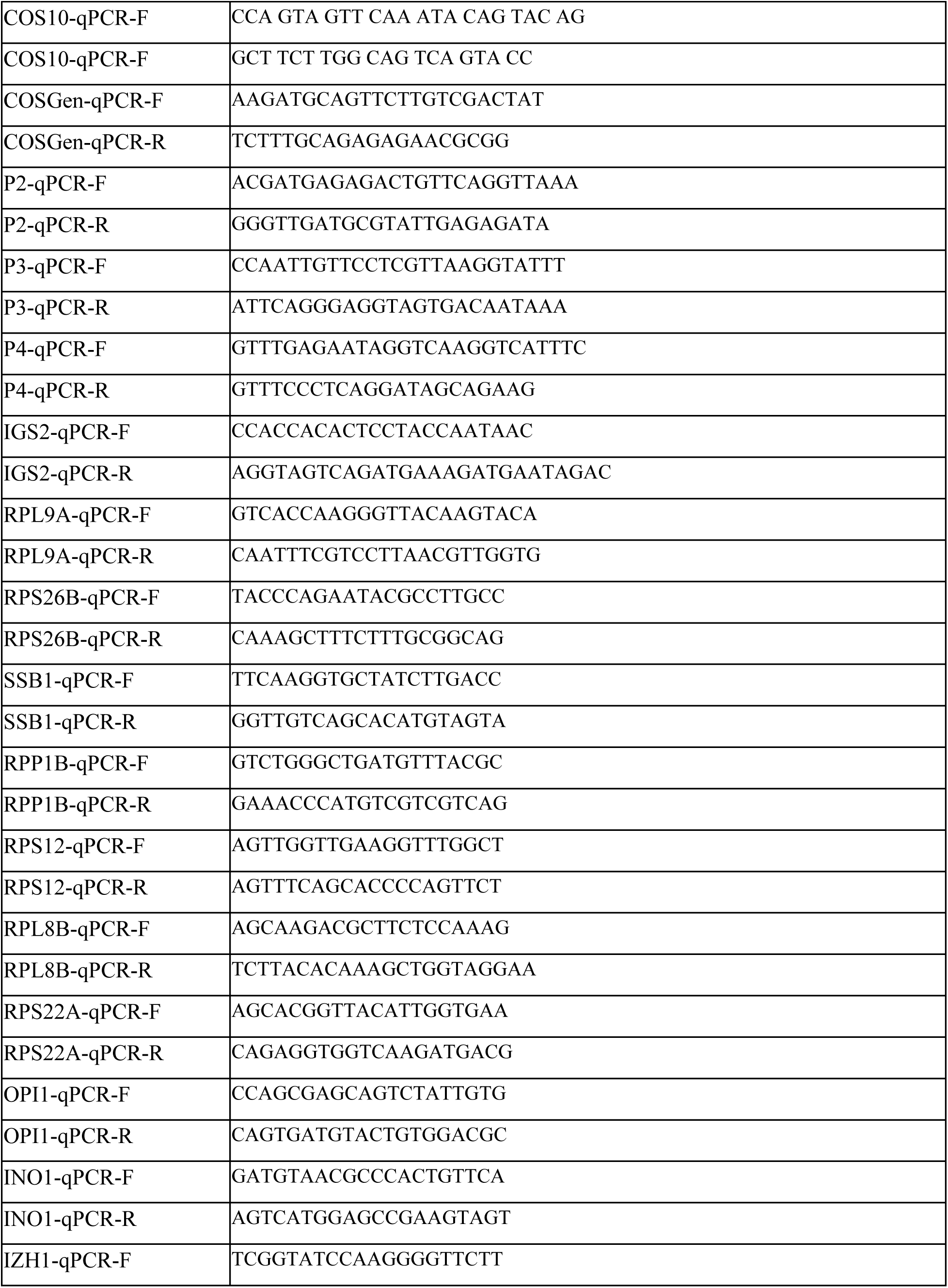
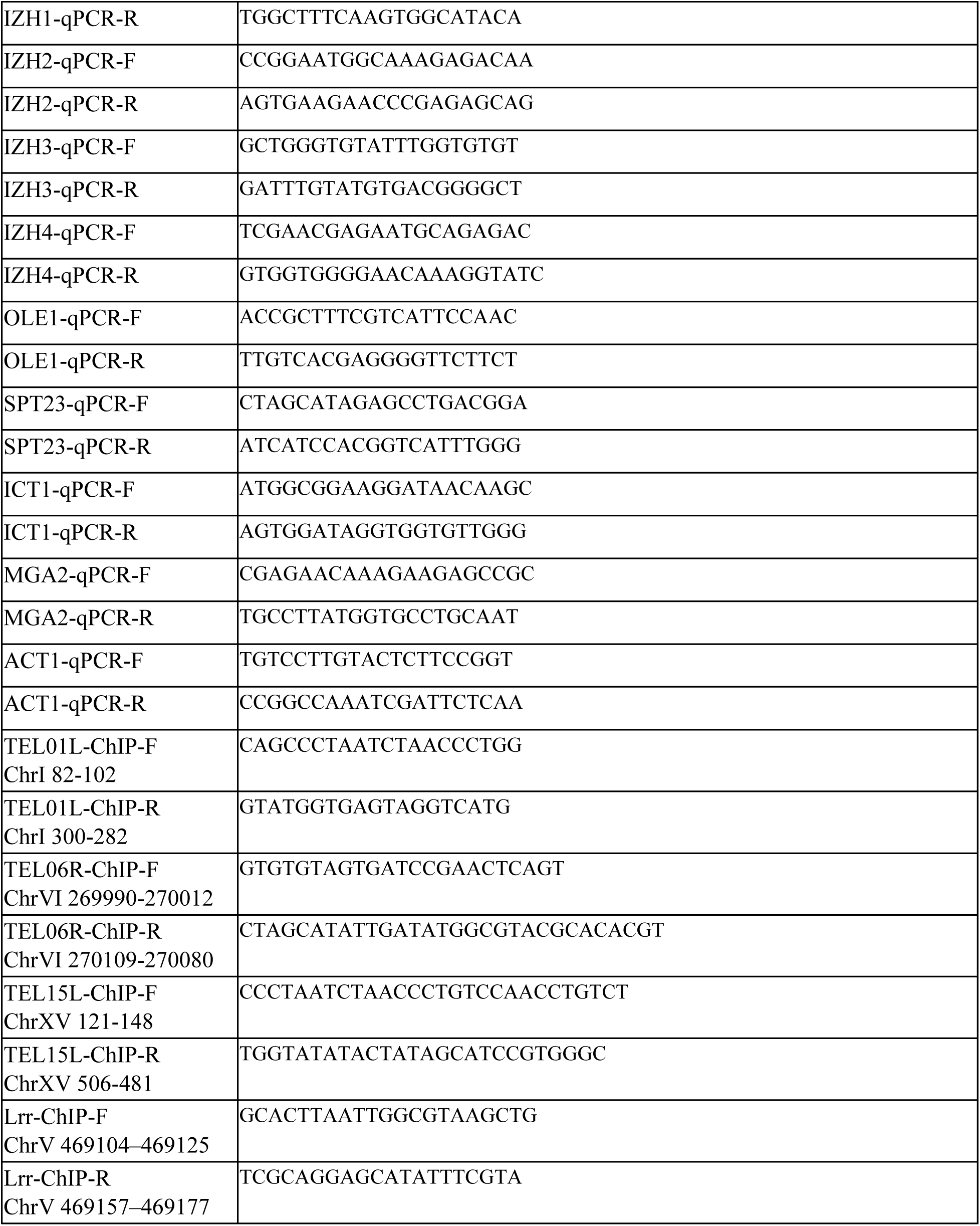
Primers used in this study

*SIR4*, *MGA2* and *SPT23* coding sequences were PCR amplified from yeast genomic DNA (lab strain W303) using gateway compatible primers (Table 5). The amplified product was then cloned into Gateway™ pDONR™221 donor vector (ThermoFisher) to generate the corresponding entry clones. Entry clones for each indicated gene were confirmed by standard DNA sequencing (DNA sequencing facility, University of Calgary). *SIR4* was then subcloned into the *S. cerevisiae* Advanced Gateway™ Destination Vector pAG415GPD-ccdB, while *MGA2* and *SPT23* were subcloned into *S. cerevisiae* Advanced Gateway™ Destination Vector pAG415GPD-EGFP-ccdB (Alberti et al., 2007). The *S. cerevisiae* Advanced Gateway™ Destination Vectors were a gift from Susan Lindquist (Addgene kit #1000000011).

### Growth conditions

Yeast strains were grown in synthetic defined minimal medium (SD) containing 0.67% yeast nitrogen base without amino acids, 2% filtered sterilized glucose and 0.1% tryptophan, 0.1% methionine, 0.1% leucine, 0.1% lysine, 0.1% arginine, 0.1% histidine, 0.02% adenine, and 0.02% uracil to fulfill strain auxotrophies. Additional adenine (0.004%, SD+Ade) was added to prevent vacuolar fluorescence in *ade2* strains used for live microscopy. Yeast cultures were routinely grown at 30 ℃ with shaking (200 rpm). Standard transformation protocol was used and cells carrying plasmids of interest were selected on selective media (Guthrie and Fink, 1991).

All experiments were conducted in exponential growth phase. Cells were routinely grown overnight in SD+Ade unless otherwise indicated. The absorbance at 600-nm (A_600_) of the cultures was monitored with a ThermoScientific™ GENESYS™ 30 Visible Spectrophotometer. Cultures were diluted the following day in fresh media and cells were grown into log phase for ∼4 hours. For experiments with edelfosine, the lyophilized drug was dissolved in ethanol as a 20 mM fresh stock. Log phase cells were standardized to A_600_ ⁓0.4, then incubated with the vehicle (0.095% ethanol) or 20 μM edelfosine at 30 ℃ with shaking for the indicated time in each experiment.

For experiments determining sensitivity to free fatty acids, defined minimal media was prepared, with glucose replaced by 4 mM oleic acid or 1.18 mM palmitoleic acid in the treatment plates. Fresh oleic or palmitoleic acid from a -20°C stock was dissolved in 500 μL DMSO (Sigma, D- 8418) then added into autoclaved warm medium to reach the final concentration. For control plates containing 2% glucose, DMSO was added to a 1% final concentration.

### Fluorescence Microscopy

Cells expressing the indicated fluorescent protein conjugates were grown into log phase as described above, then diluted to A_600_ ⁓0.4 and treated with 20 μM edelfosine or ethanol for the indicated time periods. Cells were pelleted at the indicated timepoints, resuspended in ∼10 μL of cleared media and mounted on agarose pads prepared from the same media in which the cells were cultured. Images were acquired with a Zeiss AxioImager Z2 upright epifluorescence microscope. ZEISS Zen blue imaging software and Zeiss plan Apochromat 100x/1.4 oil immersion objective lens were used for image acquisition. Z-stacks were taken in seventeen 0.24 μm steps and deconvolved using the constrained iterative algorithm available in the Zen 2.3 Pro software. Colibri 7 LED light and 90 High Efficiency filter sets were used for excitation of CFP, GFP, and DsRed. For CFP signal, samples were excited at 423/44 nm and emission was captured at 475 nm, while for GFP signal, samples were excited at 469/38 nm and emission was captured at 509 nm, and for DsRed signal, samples were excited at 555/30 nm and emission was captured at 610 nm. For images using both CFP and GFP linear unmixing was performed using the built-in settings from Zen 2.3 Pro software. Image analysis was performed manually using FIJI (Schindelin et al., 2012).

For microscopy using BODIPY™ 493/503, 1 mL aliquots of cells that had been treated with edelfosine or the vehicle for 90 minutes were incubated with 1 μL of 5 mM BODIPY in DMSO for 5 minutes at 30 ᵒC. Cells were then washed with 500 μL PBS and BODIPY signal was captured in the GFP channel. All quantitation of phenotypes was done by manual analysis of Z-stacks in FIJI, with the exception of nucleolar volume. For nucleolar volume quantification, three dimensional (X, Y, Z) stacks of yeast cells carrying Nop1^CFP^ were acquired using the CFP channel. After image deconvolution, the 3D Object Counter Volume plug-in of FIJI (Bolte and Cordelières, 2006) was used to measure the volume of the nucleolus (CFP signal) in μm^3^.

### Chromatin Immunoprecipitation

Cells expressing Sir4^MYC^ or Rap1^MYC^ were grown into log phase as described above, then diluted to A_600_ ⁓ 0.4 in 50 mL liquid media and treated with 20 μM edelfosine or ethanol for 60 minutes before crosslinking with 1.1% Formaldehyde Solution (Sigma) for 15 minutes followed by quenching with 0.125 M glycine for 5 minutes at room temperature. ChIP of yeast cells was performed as described in Lee and Keung, 2018. Briefly, cells were digested with 0.5 mg/mL zymolyase at 30 ᵒC until spheroblasts were detected using a light microscope. Spheroblasts were lysed in NP-S buffer [0.5 mM spermidine, 0.075 % NP-40, 10 mM Tris-HCl pH 7.4, 50 mM NaCl, 5 mM MgCl2, 1 mM CaCl2, 1 mM 2-mercaptoethanol, supplemented with protease inhibitor cocktail (Roche), 0.5% (w/v) Sodium Deoxycholate and 1mM PMSF] for 10 minutes on ice. DNA was sheared using a Bioruptor sonicator for four sets of 15 × 30s at amplitude of 50% with 30s shut off intervals and immunoprecipitated using sheep anti-mouse conjugated Dynabeads (Invitrogen) coupled with 1:10 mouse anti MYC antibody (Abcam, 9E10 ab32) for 2 hrs at 4°C. Immunoprecipitates were washed once with NP-S buffer and twice with wash buffer [100 mM Tris pH 8, 0.5% NP-40, 1 mM EDTA, 500 mM NaCl, 500 mM LiCl, supplemented with 0.5% (w/v) Sodium Deoxycholate, 1mM PMSF and protease inhibitor pellet (Roche)] at 4°C, each for 5 minutes with shaking at 2,200 g. Real-time qPCR reactions were carried out in technical triplicates using SYBR Green with a QuantStudio™ 6 Flex Real-Time PCR System (Applied Biosystems, Life Technologies Inc.). Ct (cycle threshold) values of Ab-coupled beads and uncoupled beads were used to calculate the fold enrichment of protein at three native sub-telomeres (Tel01L, Tel06R and Tel15L) compared to a control late replicating region on Chromosome V (469104–469177).

### Flow Cytometry

Wild-type (JC470) cells were grown into log phase as described above, then diluted to A_600_ ⁓0.4 in 10 mL liquid media. The 0-minute timepoint was taken at this time, by fixing 300 μL of culture in 1 mL 95% EtOH. Cells were then treated with either 0.095 % ethanol or 20 μM edelfosine and aliquots were fixed at 30, 60 and 90 min. Cells were stored at 4ᵒC for at least one overnight. Samples were prepared for flow cytometry by pelleting and washing with 1 mL 50 mM sodium citrate pH 7.0 solution. Cells were resuspended in 500 μL sodium citrate solution with 50 μg/mL RNAse A (ThermoFisher, EN0531) and incubated for one hour at 50 ᵒC. Samples were subsequently incubated for an additional hour at 50 ᵒC with 20 μL of 10mg/mL proteinase K (Invitrogen, AM2542). Samples were then pelleted and resuspended in 500 μL sodium citrate solution with 10 μg/mL propidium iodide (Sigma, 537059-50MG). Cell clumps were broken up by brief (3 s) sonication at 10% amplitude using a probe sonicator. Cells were then analyzed for DNA content by propidium iodide staining using a ThermoFisher AttuneNxt Flow Cytometer. PI was excited at 488 nm. The flow cytometry results were analyzed using FlowJo™ v10.8 Software (BD Life Sciences) and figures were prepared by plotting PI signal vs cell count (Becton, 2021).

### RNA-sequencing

Wild-type cells (two W303 replicates, one BY4741 replicate) were grown into log phase as described above, then diluted to A_600_ ⁓0.4 in 20 mL liquid media and treated with 20 μM edelfosine or ethanol for 60 minutes before flash-freezing. RNA was then extracted using the hot acid phenol protocol as previously described (Green and Sambrook, 2021). Briefly, yeast pellets were resuspended in 0.5 mL cold AE buffer (50mM NaOAc pH 5.2, 10mM EDTA) and lysed at 65 ᵒC by vortexing in 1.5% SDS and acid phenol. RNA and proteins were then separated with chloroform and RNA was precipitated from the supernatant using glycogen and ethanol.

RNA-sequencing and subsequent statistical analysis was performed by the Cumming School of Medicine’s Centre for Health Genomics and Informatics, University of Calgary. All samples were found to have a RIN >8 as assessed via Agilent TapeStation assay. Fluorometric assays were used to obtain the concentration in ng/μl. One μg of total RNA input was used for all libraries. NEBNext® Poly(A) mRNA Magnetic Isolation Module (E7490) and NEBNext® Ultra™ II RNA Library Prep Kit for Illumina® (E7770, E7775) were used to prepare the libraries according to the manufacturer’s instructions. Final libraries were again assessed via Agilent TapeStation assay, Qubit to obtain the concentration and Kapa qPCR to obtain the exact molarity of Illumina adapter ligated fragments according to the manufacturer’s instructions. Sequencing was then performed on an Illumina NextSeq500 system using 150 cycle mid-output 2x75 bp paired end sequencing.

Gene Ontology analysis was performed using the ClueGo plug-in for Cytoscape (Shannon et al., 2003; Bindea et al., 2009) selecting for significantly overrepresented pathways with medium network specificity and GO term fusion enabled. Transcription factor enrichment was analyzed using Yeastract (Monteiro et al., 2020), selecting for transcription factors whose targets were significantly enriched by DNA binding evidence only. Additional analysis of the RNA-sequencing data was performed with the R/Rstudio computational platform (R Core Team, 2021). Volcano plots were made with the ggplot2 package (Wickham, 2016) as a scatter plot of ln(FC) vs - log_10_(FDR). Venn diagrams were made with the venndir package (jmw86069, 2020). Telomere enrichment was calculated using the hypergeometric distribution function in base R. Principal component analysis, which concluded that genetic background minimally contributed to differences in gene expression, and it was therefore sound to average W303 and BY4741 (Figure S4A), was performed using base R functions and scores were plotted with the ggplot2 package.

*qPCR.* Wild-type or *sir4*Δ cells (W303 background) were grown into log phase as described above, then diluted to A_600_ ⁓ 0.4 in 50 mL liquid media and treated with 20 μM edelfosine or ethanol for 60 minutes before flash-freezing. RNA was then extracted using the RNeasy Mini Kit (Qiagen) as per manufacturer directions from three biological replicates. Complementary DNA (cDNA) was prepared using QuantiTect Reverse Transcription Kit (Qiagen) as indicated by the manufacturer. RNA abundance was quantified using real-time qPCR reactions, which were carried out using SYBR Green (Applied Biosystems) with a QuantStudio™ 6 Flex Real-Time PCR System (Applied Biosystems, Life Technologies Inc.). Ct (cycle threshold) values of target genes were used to calculate the fold enrichment of mRNA abundance using the ΔCt method (Livak and Schmittgen, 2001) in treated cells compared to untreated cells. Primers used are listed in Table 3.

### Spot-plating

For growth assays on solid media, cells were grown into log phase as described above and serially diluted 1:10 beginning with A_600_ ∼0.5 unless otherwise noted in the figure caption. For experiments determining sensitivity to edelfosine, cells were spotted on SD+Ade plates with 2% glucose and either 0.095% ethanol or 20 µM edelfosine using a bolt replicator and incubated at 30°C for the indicated number of days.

For experiments determining sensitivity to free fatty acids, cells were spotted on control SD + 1% DMSO plates with 2% glucose or SD + 1% DMSO containing either 4 mM oleic acid or 1.18 mM palmitoleic acid.

Images were obtained using a GelDoc imager (BioRad) and are representative plates from a minimum of three independent experiments with two technical replicates per experiment.

### Subcellular Fractionation

Membrane fractions were prepared as described previously (Marr et al., 2012). Briefly, cells were lysed using glass beads in 0.7 mL of GTE buffer [20% glycerol, 50 mM Tris-HCl, pH 7.4, 1 mM EDTA, Complete EDTA-free protease inhibitor mixture (Roche), 1 mM PMSF, 3 μg/mL pepstatin, and 1 mM phosphatase inhibitor mix (Sigma)]. Lysis was done by vortexing samples with glass beads five times for 30 s with 30 s intervals on ice in-between. Beads were washed with 0.5 mL of GTE buffer and the combined lysate was centrifuged at 16,000 × g for 15 min at 4 °C. The supernatant was then centrifuged at 450,000 × g for 15 min at 4 °C in a Beckman fixed angle rotor ultracentrifuge. Membranes were homogenized in GTE buffer using a Dounce homogenizer. Protein concentration was determined using the BCA assay (Thermo Scientific) with bovine serum albumin as a standard. Lysates and fractions (40 μg) were then analyzed by Western blot.

### Western Blot

Ten A_600_ units of cells were grown under the indicated conditions, harvested, washed once with water, and flash-frozen before being resuspended in 1 mL cold 20% trichloroacetic acid (TCA) solution. Acid-washed glass beads were added to each sample and cells were disrupted using a Mini BeadBeater (BioSpec Products) twice for 1 min each at 4 °C. The liquid was collected and combined with cold 400 uL of 5% TCA to finalize protein precipitation. Samples were then centrifuged at 3,000 rpm in a pre-cooled microcentrifuge (Eppendorf 5415R) for 10 min at 4 °C. Pellets were resuspended in 150 μL of 1x Laemmli buffer [0.2 M Tris–HCl (pH 6.8), 8% SDS, 0.4% bromophenol blue, and 40% glycerol], neutralized with 50 μL of 2M TRIS and then boiled for 5 minutes. Samples were centrifuged again for 10 min at 3,000 rpm, 40 μL of sample was analyzed by SDS-PAGE.

To capture ^GFP^Spt23, ten A_600_ units of W303 cells transformed with pAG415-EGFP-Spt23 were collected as above, but rather than being frozen, cells were immediately lysed using glass beads (1:1 volume) in 0.3 mL of GTE buffer [20% glycerol, 50 mM Tris-HCl, pH 7.4, 1 mM EDTA], supplemented with Complete EDTA-free Protease Inhibitor mixture (Roche), 1 mM PMSF and 3 μg/mL pepstatin. Lysis was done as described above using glass beads. Samples were centrifuged at 4000 rpm × 5 min at 4°C in a pre-cooled microcentrifuge (Eppendorf 5415R) and the supernatant was collected. After lysis, 10 μl of 4X SDS was added to 30 μl of samples, which were incubated for 2 minutes at room temperature before loading into the gel.

For western blot analysis, proteins were separated by 8% resolving gel containing trichloroethanol (TCE; Sigma) to visualize proteins (Ladner et al., 2004). Proteins were transferred to a polyvinylidene fluoride (PVDF) membrane (Millipore) using a Bio-Rad transfer system at 100 V for 1 h or 25 V for 16 hrs, and then stained with Red Ponceau (Sigma) to confirm transfer. The following primary antibodies were used: 1:5,000 mouse αMYC (Abcam, 9E10 ab32), 1:1,000 mouse αRap1 (Santa Cruz, sc374297), 1:1,000 mouse αGFP (Roche, 11814460001), 1:500 mouse αPgk1 (Molecular Probes), 1:1,000 rabbit αH2A (Active Motif, 39235), 1:1,000 rabbit αH2A^S129ph^ (Active Motif, 39271) and 1:2,000 mouse αRad53 (Abcam, ab104232). Horseradish peroxidase conjugated secondary antibodies (Invitrogen, Thermo Fisher) and enhanced chemiluminescence (Amersham, GE Healthcare) were used for detection using an Invitrogen iBright FL1500 Imaging System.

### Lipid extraction and neutral lipids analysis

Lipid extracts were prepared as described previously (Zaremberg and McMaster, 2002). Briefly, cells were concentrated by centrifugation, washed twice with water, and resuspended in 1 mL of CHCl_3_/CH_3_OH (1/1, v/v), then stored overnight at - 20 ᵒC. Cells were disrupted for 1 minute at 4 °C in a Mini BeadBeater (BioSpec Products) with acid-washed glass beads. Beads were washed with 1.5 mL of CHCl_3_/CH_3_OH (2/1, v/v), then 0.5 mL of CHCl_3_ were added to the combined supernatant followed by 1.5 mL of water to facilitate phase separation. Lipids were dried under nitrogen, resuspended in 30 μL chloroform and 15 μL of this resuspension was loaded for separation on silica gel on TLC aluminum foils (Sigma-Aldrich, 60805-25EA) using a solvent system containing 80:20:1 petroleum ether/diethyl ether/acetic acid for separation of neutral lipids. Standard mix (10 μg/ each lipid) was loaded onto the plate. Standard mix consisted of a mix of 2 mg/mL solution of DAG, ergosterol, oleic acid, TAG, and cholesteryl oleate. Plates were developed with iodine vapors and imaged using a BioRad GelDoc. Images shown are representative of two independent experiments with two technical replicates per experiment.

Free fatty acid levels were calculated using densitometry, by measuring the pixel intensity of sample bands and comparing to a standard curve of 0.25, 0.5, 1 and 2 μg of pure oleic acid. Samples were normalized by determining total phosphate released from phospholipids. Phosphate was determined using the AMES assay (Ames, 1966). Briefly, lipid extracts were resuspended in chloroform and dried in glass test tubes. A standard curve of 0, 0.5, 1, 2, 4 and 6 mM NaH_2_PO_4_ solution was prepared alongside the samples and 30 μL of 10% w/v magnesium nitrate in ethanol was added to all tubes and ashed using a flame. The ash was then resuspended in 300 μL 0.5M HCl and boiled for 15 minutes. The free inorganic phosphate from this reaction was then visualized by adding 700 μL 1:6 10% w/v ascorbic acid in water : 0.42% w/v ammonium molybdate in 0.5M H_2_SO_4_ and developed for one hour at 37ᵒC. All samples were prepared in duplicate. Absorbance was measured at 820 nm using a Biotek Synergy H1 Microplate Reader.

### Statistical analysis

All statistical analysis was performed in GraphPad Prism 9.3.1. The following analyses were conducted depending on the experiment as indicated in each figure legend. A minimum of three independent experiments were analyzed in each case.

For categorical quantification analyzing the difference between only two variables, two-tailed unpaired t-tests were performed. For analyzing the difference between more than two variables, standard one-way ANOVA with Tukey’s multiple comparisons post-test was used. For multiple comparisons, multiple unpaired t-tests were performed and corrected for multiple comparisons using the Holm-Šídák post-test (Holm, 1979). Additionally the Welch’s correction for unequal variance (Welch, 1947) was used when appropriate. For analysis of differences between wildtype and mutants in treated and untreated conditions, 2-way ANOVA with Šídák’s multiple comparisons test was applied (Sidak, 1967).

For numerical quantification analyzing the difference between only two variables, two-tailed nested t-tests were performed. For analysis of differences between wildtype and mutants in treated and untreated conditions, nested one-way ANOVA with Šídák’s multiple comparisons pos-test were performed.

## Supplementary Data

**Figure S1** is related to Figure 1, checking for DNA damage response in edelfosine.

**Figure S2** is related to Figure 2, providing additional microscopy images and telomere interactions with the nuclear membrane, as well as cell cycle effects of edelfosine.

**Figure S3** is related to Figure 3, with protein and sensitivity controls for Sir4 as well as qPCR gene expression of sub-telomeric genes.

**Figure S4** is related to Figure 4, including principal component analysis and gene ontology of the RNA-seq, as well as qPCR of rDNA.

**Figure S5** is related to Figure 6, providing additional microscopy images of the ER.

## Supplementary Tables

**Table S1** lists all genes differentially expressed in edelfosine.

**Table S2** lists all sub-telomeric genes found to be significantly differentially expressed in edelfosine with chromosome coordinates.

**Table S3** lists known internal gene targets of the SIR complex downregulated in edelfosine. **Table S4** lists known ribosomal protein genes bound by Rap1 downregulated in edelfosine. **Table S5** lists all lipid metabolism related genes differentially regulated in edelfosine.

---

**Supplementary Figure 1.**
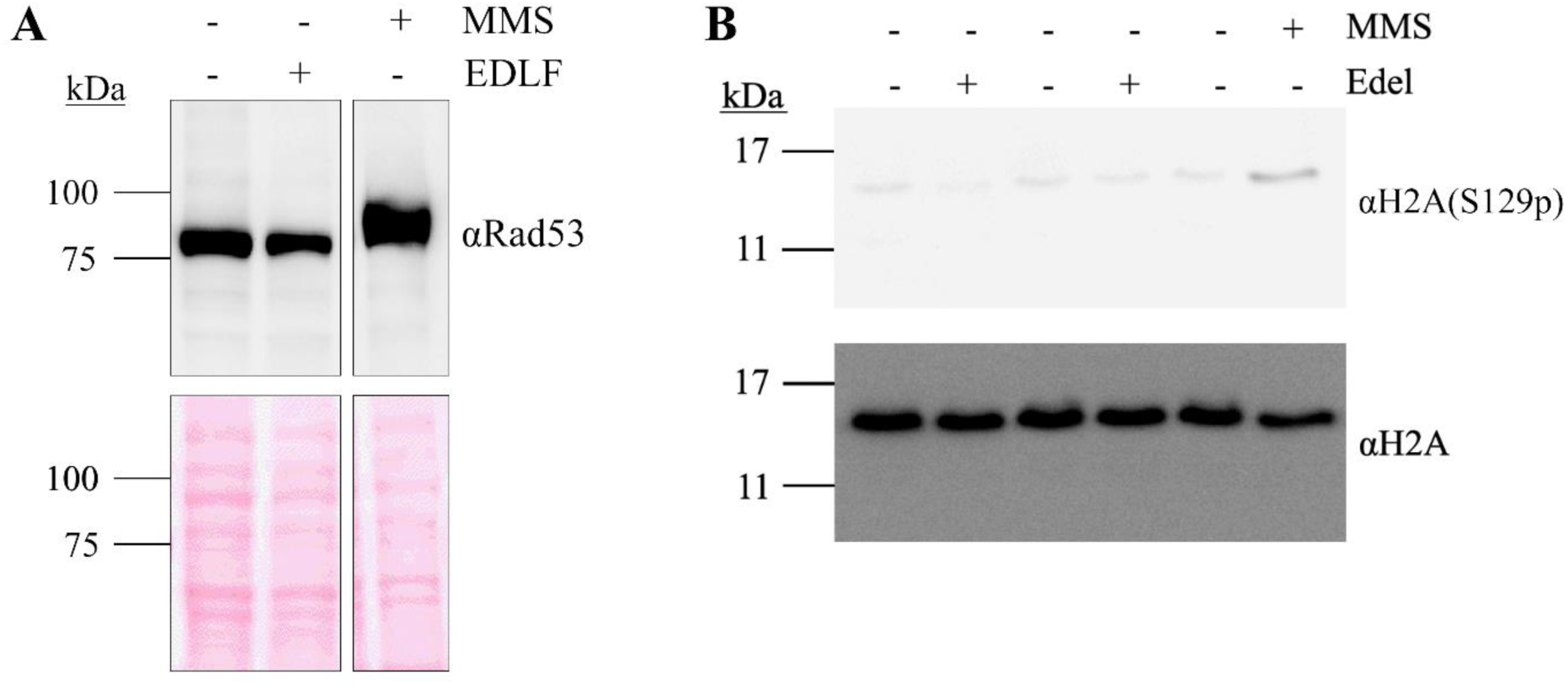
Edelfosine does not elicit a DNA damage response. (**A**) Anti-Rad53 western blot of whole cell lysates from cells (W303) treated with edelfosine, vehicle or 0.01% MMS for 60 minutes. Bottom panel shows protein loading by Red Ponceau. (**B**) Anti-phospho-Ser^129^-H2A and anti-total H2A blot of whole cell lysate from cells expressing Sir4-13MYC or Heh1-13MYC treated with edelfosine, the vehicle or 0.01% MMS for 60 minutes.

**Supplementary Figure 2.**
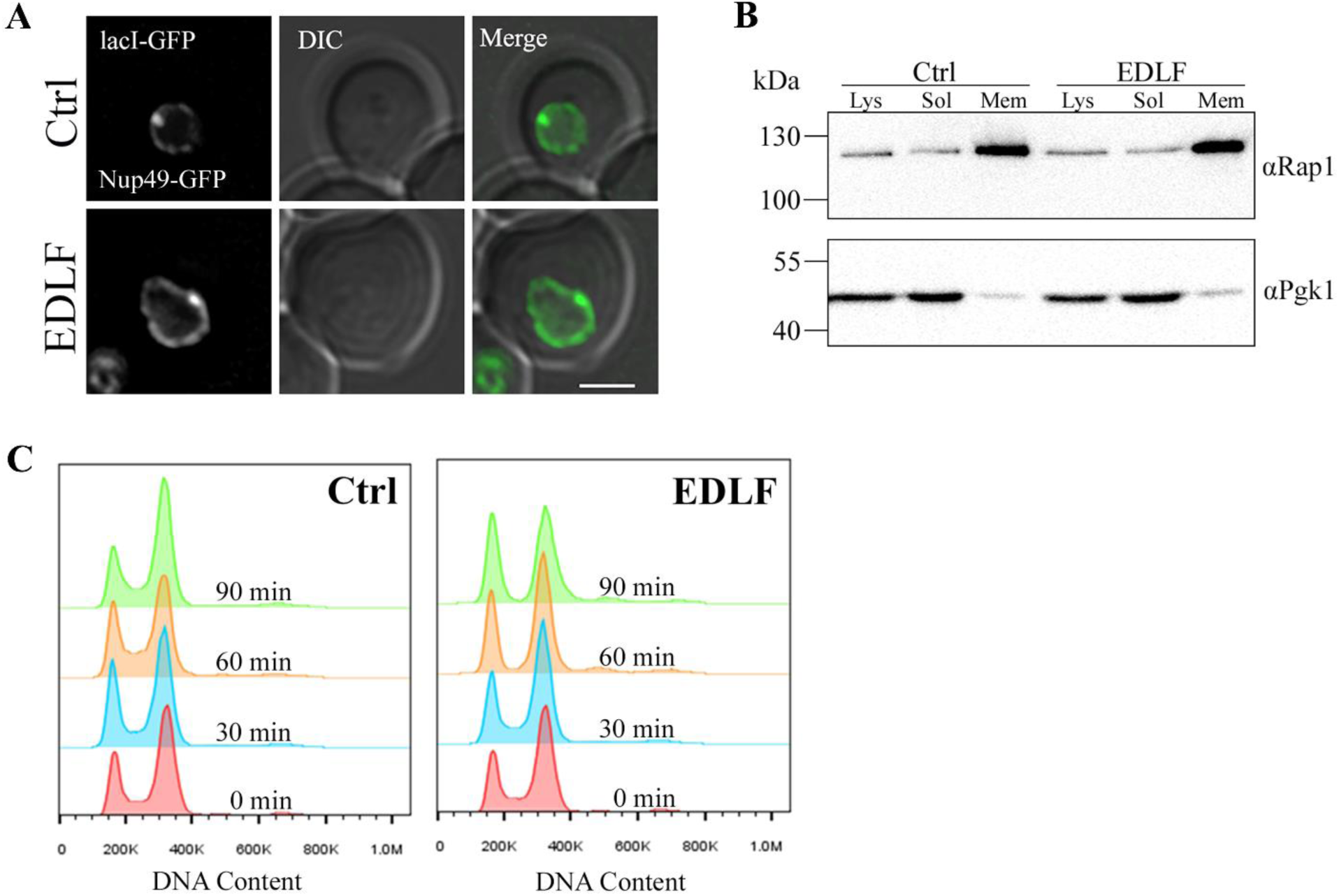
Telomere interactions with the nuclear membrane are preserved in edelfosine. (**A**) Representative cells expressing the Tel06R constructs quantified in Figure 2B were imaged using live fluorescence microscopy after 90 minutes in edelfosine or the vehicle. Scale bar represents 2 µm. (**B**) Membrane and soluble fractions were collected from BY4741 cells expressing Spt23-TAP treated with 20 μM edelfosine or vehicle for 60 minutes as described in Materials and Methods, then blotted for detection of endogenous Rap1 and Pgk1. (**C**) Cell cycle analysis of wild type cells (W303) treated with edelfosine or vehicle for the indicated times. For all panels: Ctrl = vehicle; EDLF = edelfosine.

**Supplementary Figure 3.**
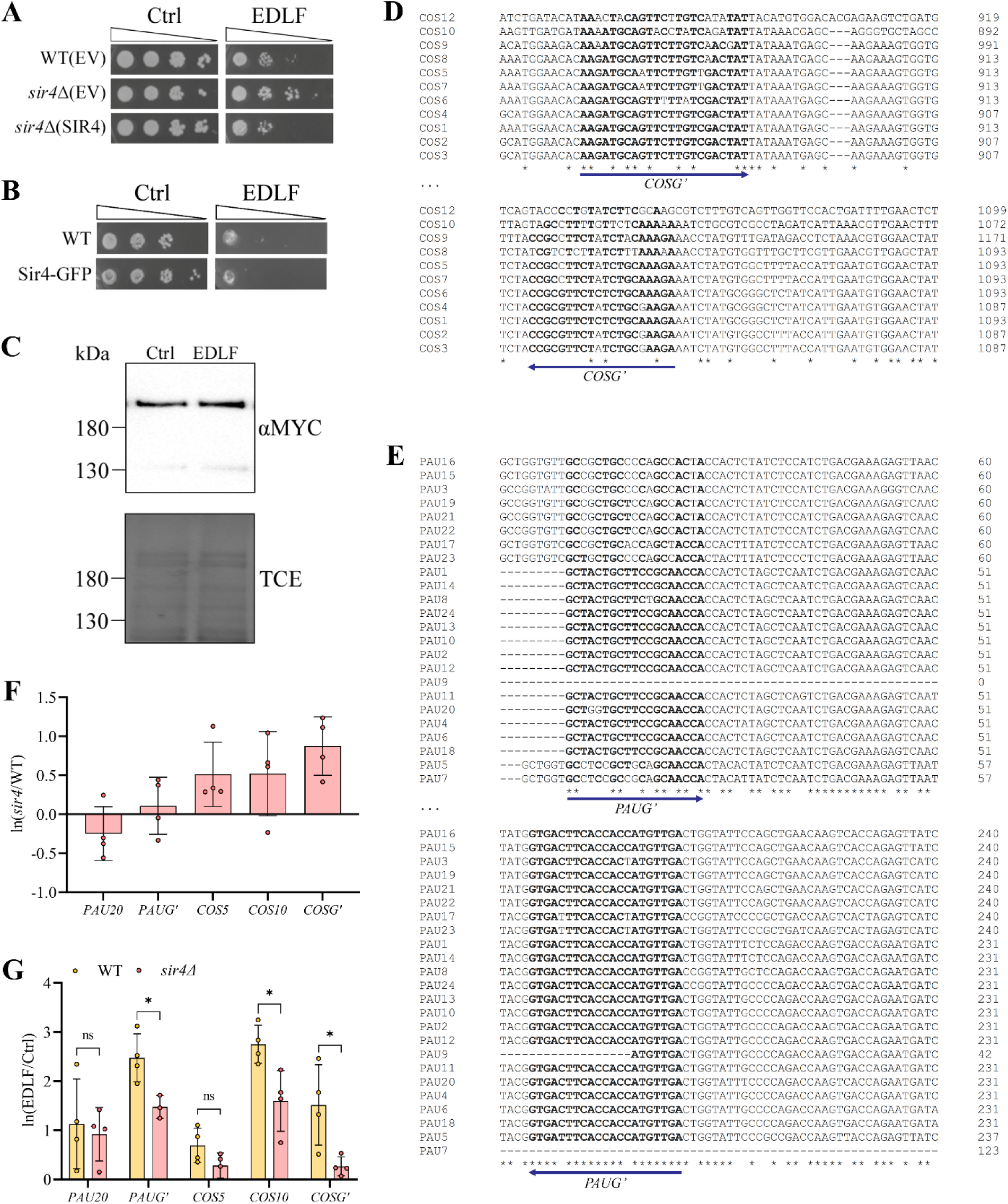
Treatment with edelfosine induces expression of the SIR-dependent COS and PAU sub-telomeric gene families. (**A)** Growth of wild-type (W303) or *sir4Δ* cells expressing *SIR4* from a centromeric plasmid or the empty vector (EV) on SD-Leu+Ade plates containing 20 μM edelfosine or vehicle. Plates were incubated at 30℃ and imaged after three days of growth. (**B**) Growth of wild-type (W303) or endogenously tagged *SIR4*-GFP cells on synthetic solid medium containing 20 μM edelfosine or vehicle. Plates were incubated at 30℃ and imaged after two days of growth. (**C**) Western blot of Sir4^MYC^ after 60 minutes with edelfosine or control. Bottom panel shows protein loading by TCE. (**D, E**) Sequence alignment of the highly similar COS (**D**) and PAU (**E**) family genes at the regions amplified by qPCR. Bold letters indicate binding sites for the general primers. (**F**) qPCR of the indicated genes in *sir4*Δ cells relative to wild type, expressed as ln(mutant/WT) or (**G**) in wild-type and *sir4*Δ cells treated for 60 minutes with edelfosine or vehicle expressed as ln(EDLF/Ctrl). Bars represent mean ± SD for three independent experiments while circles represent individual experiments. * indicates p-value <0.05 as determined by unpaired t tests with Holm-Šídák correction for multiple comparisons. For all panels: Ctrl = vehicle; EDLF = edelfosine.

**Supplementary Figure 4.**
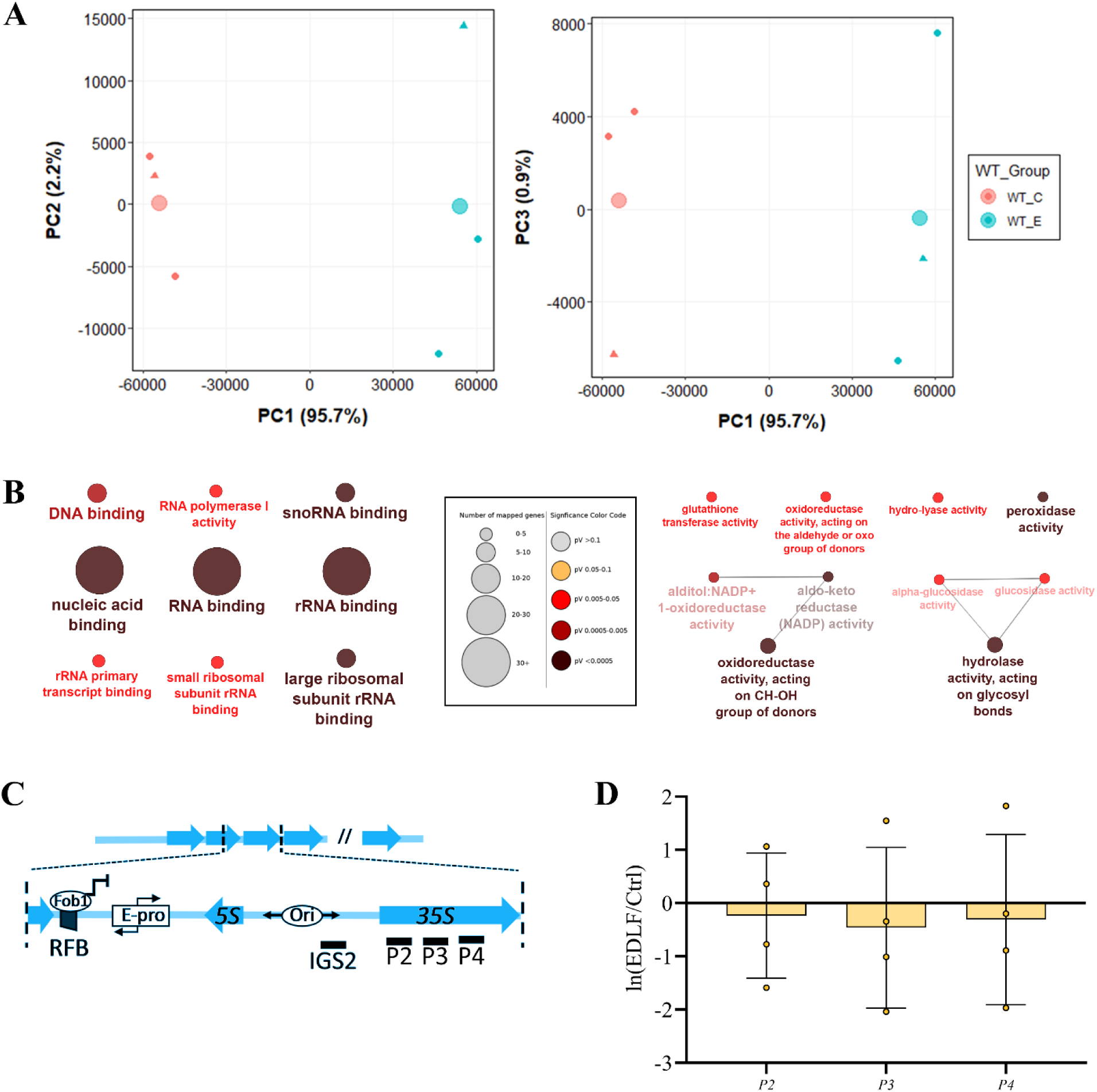
Genetic background and gene ontology analysis of RNA- sequencing hits. (**A**) Scatter plots of the first three principal component analysis scores for the transcriptional landscape of each sample submitted for RNA-sequencing. W303 samples are represented as circles while BY4741 samples are represented as triangles. Large circles indicate the mean PC score for each treatment. Red is Ctrl and blue is EDLF. (**B**) Molecular function GO terms enriched in genes downregulated (left) and upregulated (right) in edelfosine more than ln(1.5-fold). Legend demonstrating correlation of size and colour with the number of genes per term and the significance score, respectively, is in the middle. (**C**) Schematic of rDNA repeats showing regions amplified by qPCR primers. (**D**) qPCR of 35S rDNA targets (P2-4) in WT cells treated for 60 minutes with edelfosine or vehicle expressed as ln(EDLF/Ctrl) and normalized to *IGS2*. Bars represent mean ± SD for three independent experiments while circles represent individual experiments. For all panels: Ctrl = vehicle; EDLF = edelfosine.

**Supplementary Figure 5.**
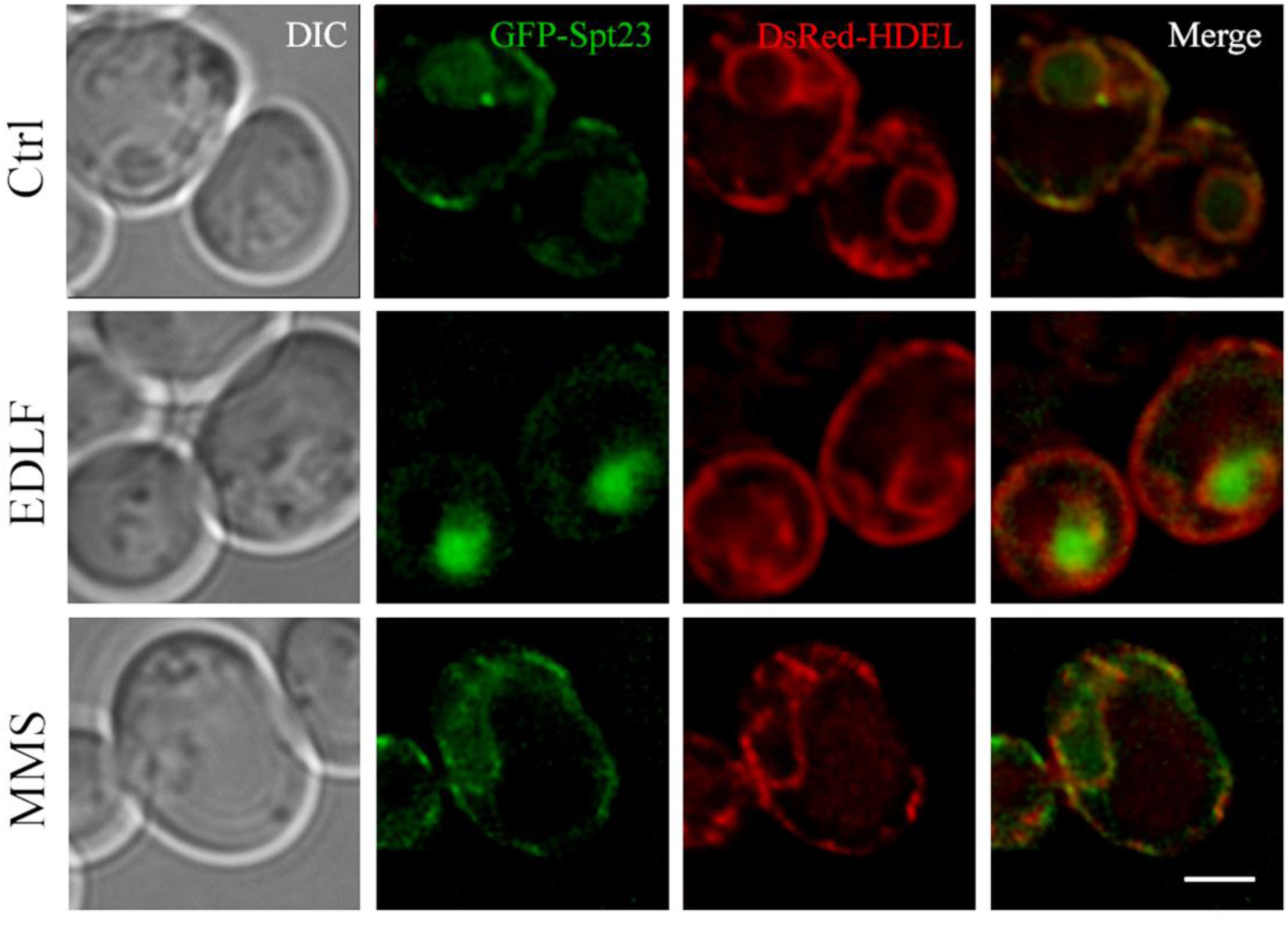
Spt23 translocates to the nucleus in response to edelfosine. Representative images of wild-type (W303) cells expressing ^GFP^Spt23 from a centromeric plasmid under the constitutive GPD promoter and ^DsRed^HDEL that were imaged after growth in SD- Leu+Ade and 60 minutes in the presence of edelfosine (EDLF), the vehicle (Ctrl), or 0.1% methyl methane sulfonate (MMS).

## Acknowledgments

The authors would like to thank CM Unger and G Bertolesi for their helpful discussions, S. Gasser for sharing microarray data of internal SIR targets and S Gasser and S Emr for the kind gift of strains and plasmids. This work was supported by a Discovery Grant and a Discovery Accelerator Supplement from the Natural Sciences and Engineering Research Council (NSERC) to VZ and by operating grants from CIHR MOP-82736; MOP-137062 and NSERC 418122 awarded to JAC. MLSP is funded by a doctoral fellowship from NSERC and Alberta Innovates.

